# Molecular dynamics simulations reveal a mechanism of calcium homeostasis driving homotrimerisation and heterotrimerization of type I collagen

**DOI:** 10.1101/2025.04.03.646381

**Authors:** Emily J Johnson, João V de Souza, Agnieszka K Bronowska, Elizabeth G Canty-Laird

## Abstract

Type I collagen is the main structural protein of vertebrates and forms molecular trimers from the COL1A1 and COL1A2 gene products; pro-alpha-1(I) and pro-alpha-2(I)), during biosynthesis. The amino acid sequence of the C-propeptide of collagen, which is removed before collagen fibril formation, initially drives heterotrimerisation and calcium ions are required for trimers to form. The homotrimeric form is associated with age-related diseases including cancer, fibrosis, musculoskeletal and cardiovascular conditions but circumstances under which the abnormal homotrimer may form were poorly understood. Here we used molecular dynamics simulations of the C-propeptide protein structure, to show that intra- and intra-chain hydrogen bonding is affected by the loss of calcium and that chains become destablised, particularly at the interfaces of each chain. Loss of calcium resulted in an increased distances between cysteine residues that form inter-chain disulphide bonds, predicting an inability for disulphides to form in the absence of calcium. Pulling simulations and modeling calcium dissociation from monomers showed that calcium ions were more strongly bound to the alpha-1(I) than the alpha-2(I) chain. However, pulling a single alpha chain from the heterotrimer or homotrimer demonstrated that the alpha-2(I) chain had a higher affinity to trimers than a third alpha-1(I) chain. Hence although heterotrimerisation is normally favoured, in reduced calcium conditions the homotrimer can form by sequestering available calcium to the alpha-1(I) chains. This study provides a molecular explanation for a calcium-based mechanism driving heterotrimerisation versus homotrimerisation of type I collagen.

## Introduction

Type I collagen is normally a heterotrimer composed of two α1(I) chains and one α2(I) chain, derived from the COL1A1 and COL1A2 genes respectively. N- and C-terminal globular propeptide domains flank a 300nm long right-handed triple helical domain with a pitch of 18 amino acids per turn that supercoil around a central axis. The helical region has a repeating Gly-X-Y amino acid structure, where X and Y are often proline and hydroxyproline respectively. The glycine residues are positioned at the centre of the triple helix, allowing the bulky side chains to occupy the outer positions and enabling tight packing (1). The C- and N-terminal propeptides confer solubility to the chain, preventing premature aggregation. The C-propeptide guides the trimerisation process, as chain selection and alignment begins with C-propeptide trimerisation, after which folding of the triple helix occurs from the C- to N- end (2). The propeptides are removed to facilitate assembly of trimeric type I collagen molecules into fibrils.

An abnormal homotrimeric form composed of three α1(I) chains has been reported in adult skin and embryonic tissues (3,4). However, the homotrimeric form is also associated with diseases such as cancer, osteoarthritis, osteoporosis, fibrosis and Ehlers-Danlos syndrome (5,6). Molecular dynamics simulations of a 57 amino acid region of the >1000 amino acid triple helical region have also shown the homotrimer to be softer and more flexible (7). The homotrimeric helix freely rotates and forms kinks in the Gly-X-Y domain which is predicted to lead to greater lateral distances between homotrimeric molecules in fibrils, consistent with experimental findings (8). Altered packing may be responsible for reported differences in intermolecular collagen crosslinking in the osteogenesis imperfecta murine (oim) mouse model (9-11), although the homotrimeric collagen is itself not responsible for bone fragility (12). Type I collagen homotrimer is resistant to proteolysis compared to the heterotrimer and degradation by MMP-1 is approximately ten times slower (13). The homotrimer however appears more sensitive to degradation under mechanical strain, whilst the heterotrimer is less sensitive (14,15). As the two trimeric forms of type I collagen demonstrate such different biophysical, dynamic and structural properties, their physiological roles presumably differ, and synthesis needs to be tightly controlled to ensure the correct type of type I collagen is being produced.

The C-propeptide drives fibrillar collagen trimerisation and also determines trimer chain composition. In type III collagen, the chain recognition sequence coordinates trimerisation and ensures that only α1(III) homotrimers form (16,17). This chain recognition mechanism does not however occur in type I collagen, where interchain interactions occur at key residues that form salt-bridges (5) and trimer composition is partially governed by a network of disulphide bond-forming cysteines in the C-propeptide (18). The α1(I) chain C-propeptide contains eight cysteine residues (Cys 1-8), two of which participate in inter-chain disulphide bonding: C2 and C3. The α2(I) chain C-propeptide lacks the C2 residue and can only form one inter-chain disulphide bond. This ensures that only heterotrimers and α1(I) homotrimers can form and has been termed the ‘cysteine code’. The same study demonstrated a key role for calcium ions in mediating trimerisation, as in the absence of available calcium in solution no heterotrimers or homotrimers can form. The α1(I) and α2(I) C-propeptides contain a conserved calcium-binding loop coordinating a structural calcium ion that sits at subunit interfaces in the C-propeptide trimer. Indeed, a COL1A1 mutation substituting a calcium binding residue in the C-propeptide of the α1(I) prevents trimersation and results in perinatal lethality (18,19).

Proteins are flexible entities, and their dynamic properties can play a crucial role in their functionality. Molecular dynamics is a computational technique that has been termed a ‘molecular microscope’ as it applies the laws of physics to simulate protein dynamics over time. A requirement for molecular dynamics is a structural model of the protein in question. Though the type I collagen C-propeptide heterotrimer structure has yet to be solved, several high quality crystal structures of other collagen C-propeptides exist, including the type I collagen homotrimer (5). The α2(I) C-propeptide chain’s high sequence identity with the α1(I) C-propeptide chain means a 3D structure can be estimated using homology modelling. In the present study, equilibrium molecular dynamics simulations along with enhanced sampling techniques were used to study type I collagen C-propeptide stability, with and without the structural calcium bound, to investigate how the structural calcium guides trimerisation, and how calcium homeostasis might play a role in homotrimer production.

## Results

### The C-propeptide shows increased RMSD but decreased radius of gyration over 1000 simulation

A structure for the type I collagen heterotrimeric C-propeptide was obtained by homology modelling using the crystal structure of the homotrimeric form. Simulations were then carried out over 1000ns for both the heterotrimer (Fig. 1A) and homotrimer (Fig. 1B), either including (apo form) or excluding (holo form) the three calcium ions bound at the interfaces between sub-units. The root mean square deviation (RMSD) increased for each trimer over the course of the simulation (Fig. 1C), whilst the radius of gyration decreased (Fig. 1D).

**Figure 1:**
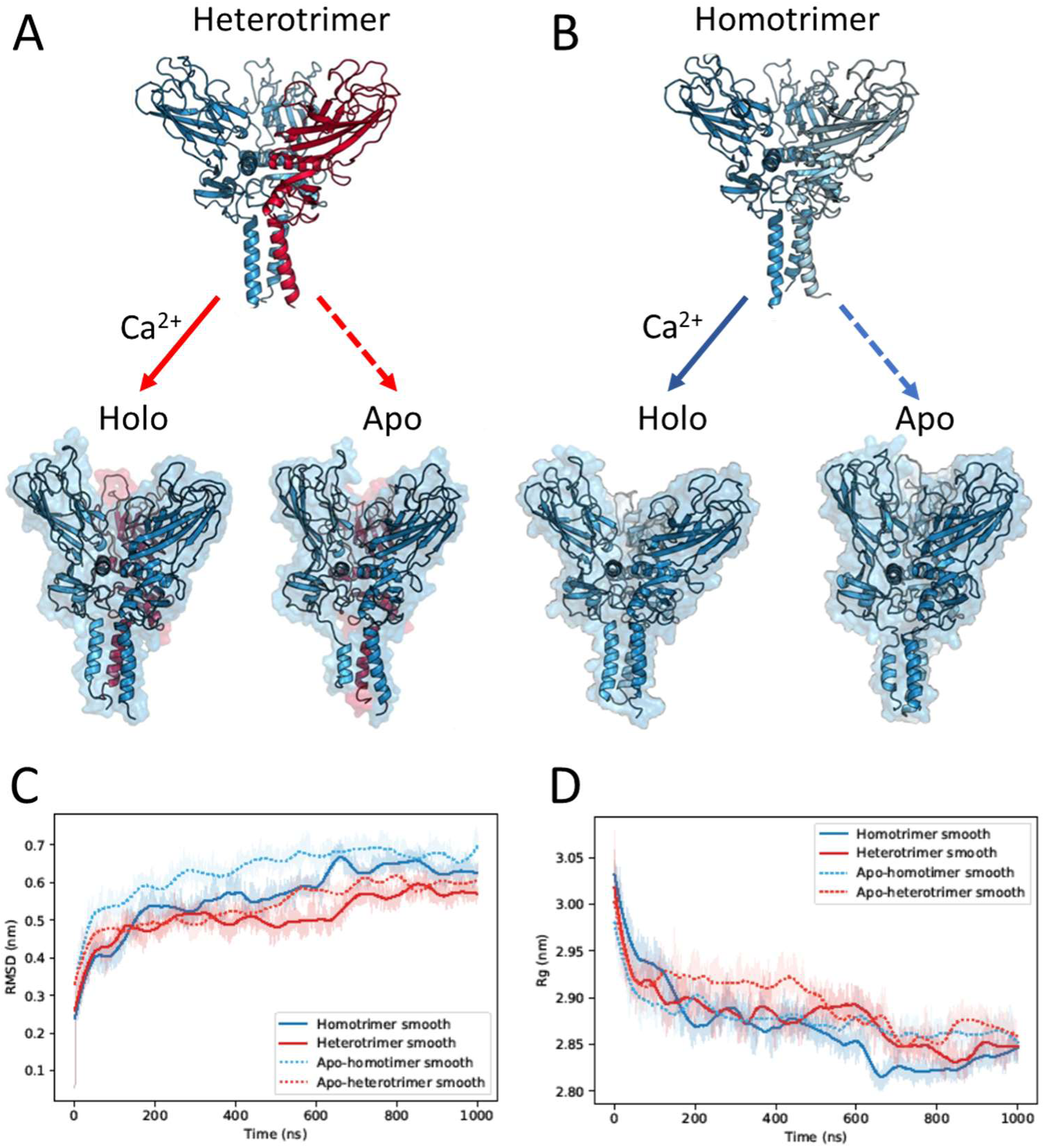
Molecular dynamics simulations of the calcium-replete and depleted C-propeptide heterotrimer and homotrimer. Starting structures of the heterotrimer (A) and homotrimer (B) compared to the holo- (replete) and apo- (depleted) forms after 1000 ns of simulation. The α2(I) chain is shown in red for the heterotrimer structures. Images are derived from one of three repeat simulations. C: Root mean square deviation (RMSD, nm) of each trimer over 1000 ns of simulation. D: Radius of gyration (Rg, nm) of each trimer over 1000 ns of simulation. For RMSD (B) and Rg (C), average values of three equilibrium simulations are shown using a transparent fill, with the smoothed values are overlaid.

Overall, the heterotrimer appeared more stable than the homotrimer with a mean RMSD of 0.511 nm versus 0.565 nm, and the holo- proteins having lower RMSDs, indicating higher stability than the corresponding apo- structures (Table 1). The homotrimer appeared to maintain a tighter fold, and hence to be more compact than the heterotrimer, based on lower Rg values. There was a statistically significant difference in time-averaged RMSD values within the trimer group (Table 2) but pairwise comparisons for linear mixed models fitted to the RMSD and Rg data did not detect significant differences between trimer types.

**Table 1:** Time averaged structural properties calculated for the homotrimer, heterotrimer, apo-homotrimer and apo-heterotrimer across three replicates. Standard error values are given in parentheses.

| Trimer type | Backbone RMSD (nm) | Backbone-Rg (nm) |
| --- | --- | --- |
| Holo-homotrimer | 0.565 (0.00043) | 2.866 (0.00022) |
| Holo-heterotrimer | 0.511 (0.0003) | 2.878 (0.00016) |
| Apo-homotrimer | 0.632 (0.00033) | 2.877 (0.00011) |
| Apo-heterotrimer | 0.540 (0.0003) | 2.895 (0.00015) |

**Table 2:**
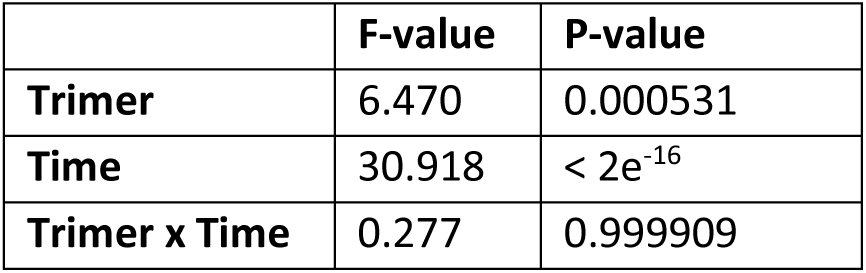
A repeated measures two-way ANOVA to assess whether there were differences between the RMSD values for trimers over time. All three replicates were included in the statistical model.

|  | F-value | P-value |
| --- | --- | --- |
| Trimer | 6.470 | 0.000531 |
| Time | 30.918 | $< 2e^{-16}$ |
| Trimer x Time | 0.277 | 0.999909 |

### Hydrogen bonding is affected by the loss of calcium

To investigate the impact of calcium on the hydrogen bonding network it was probed using the Cytoscape- Chimera StructureViz package (Figure 2, Supplementary Figure S1 and Supplementary Table 1.). For the homotrimer, residues CYS-64 (C3), ASP- 67 and ASN-61 of the calcium-binding loop form hydrogen bonds with ASP-43, ARG-42 and ARG-39 respectively on the neighbouring chain, which is an alpha-helix region containing the partner C2 for C2-C3 binding. These hydrogen bonds are conserved between the α1(I) and α2(I) chains. The α1(I) central alpha-helical region contains an ASP-129 residue that also binds ARG-42 on the neighbouring chain, forming a salt bridge (ALA-128 can also hydrogen bond ARG-42, but not as strongly), as previously described (5). In the heterotrimer, the α2(I) chain has a GLU-130 residue that forms a salt bridge with ARG-42 on the neighbouring chain instead.

**Figure 2:**
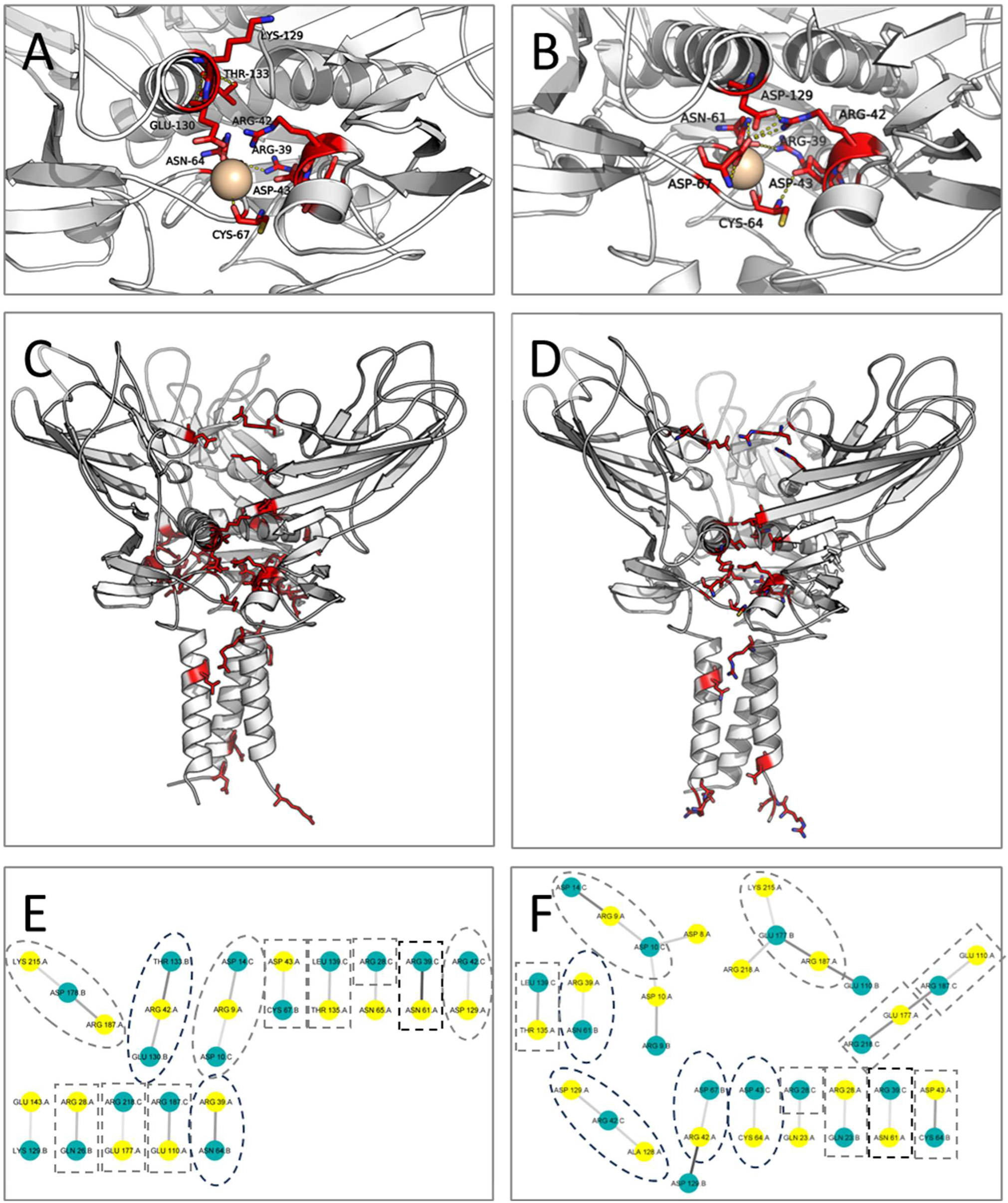
C-propeptide heterotrimer and homotrimer hydrogen bonding network revealed by StructureViz analysis. A-D: Trimer cartoons with residues that participate in inter-chain hydrogen bonds shown in red and as ‘stick’ representations. A, B: Hydrogen bonds at the chain interface for the heterotrimer (A) and homotrimer (B). C, D: Inter-chain hydrogen bonds for the whole structure for the heterotrimer (C) and homotrimer (D). The calcium ion is shown as a wheat-coloured sphere. The N-terminal regions demonstrate increased flexibility due to the lack of Gly-X-Y domain. E, F: Hydrogen bonding network for the alpha-1(1) chain from explicit hydrogen MD simulations for the heterotrimer (E) and homotrimer (F). Only one replicate was carried out with explicit hydrogens, with a time step of 2 fs. Alpha-1(1) is chain A is (yellow); alpha-1(2) in the homotrimer and alpha-2 in the heterotrimer is chain B (teal); and alpha-1(3) is chain C (also teal). The edge weight corresponds to how conserved the bond was throughout the simulation. The darker edges represent bonds that were present throughout most of the simulation, the lighter ones were more transient bonds. Black circles denote bonds referred to in the text. Grey circles denote bonds that are conserved, or similar, between the heterotrimer and homotrimer.

These inter-chain interactions are stabilised by intra-chain hydrogen bonds. For the α1(I) chains the ARG-39 and ASN-61 hydrogen bond is stabilised by ASN-61 forming a hydrogen bond with GLN-133, and ARG-39 forming hydrogen bonds with PRO-60 and GLN-62. The ARG-42 and ASP-67/ASP-129 salt bridges are stabilised by ARG-42 forming hydrogen bonds with LEU-246 and THR-142. ASP-43 forms an intra-chain salt-bridge with ARG-39, which may act to stabilise the protein structure and overall inter-chain hydrogen bonding network, as ARG-39 is one of the residues that participates in binding at the interface. In the heterotrimer, the stabilising intra-chain hydrogen bonds for the α2(I) chain are between ARG-45 and PHE-246, ASN-64 and GLN-134, and ASP-70 and GLN-134.

In addition to ASP-43 forming an inter-chain hydrogen bond with CYS-64 and intra-chain salt-bridge with ARG-39, it also forms an intra-chain hydrogen bond with CYS-47 (i.e. it binds both partners in C2-C3 disulphide bond formation). It may be that this residue shuttles between forming hydrogen bonds with one cysteine or the other, coordinating their association.

To assess whether the hydrogen-bonding networks at the chain interfaces are affected by calcium loss, root mean square fluctuation (RMSF) values were calculated per residue throughout each 1000 ns simulation and averaged across the three replicates (Figure 3). RMSF hotspots occurred around residue 100 (in a petal loop region) and residue 125 which is in the base interface region.

**Figure 3:**
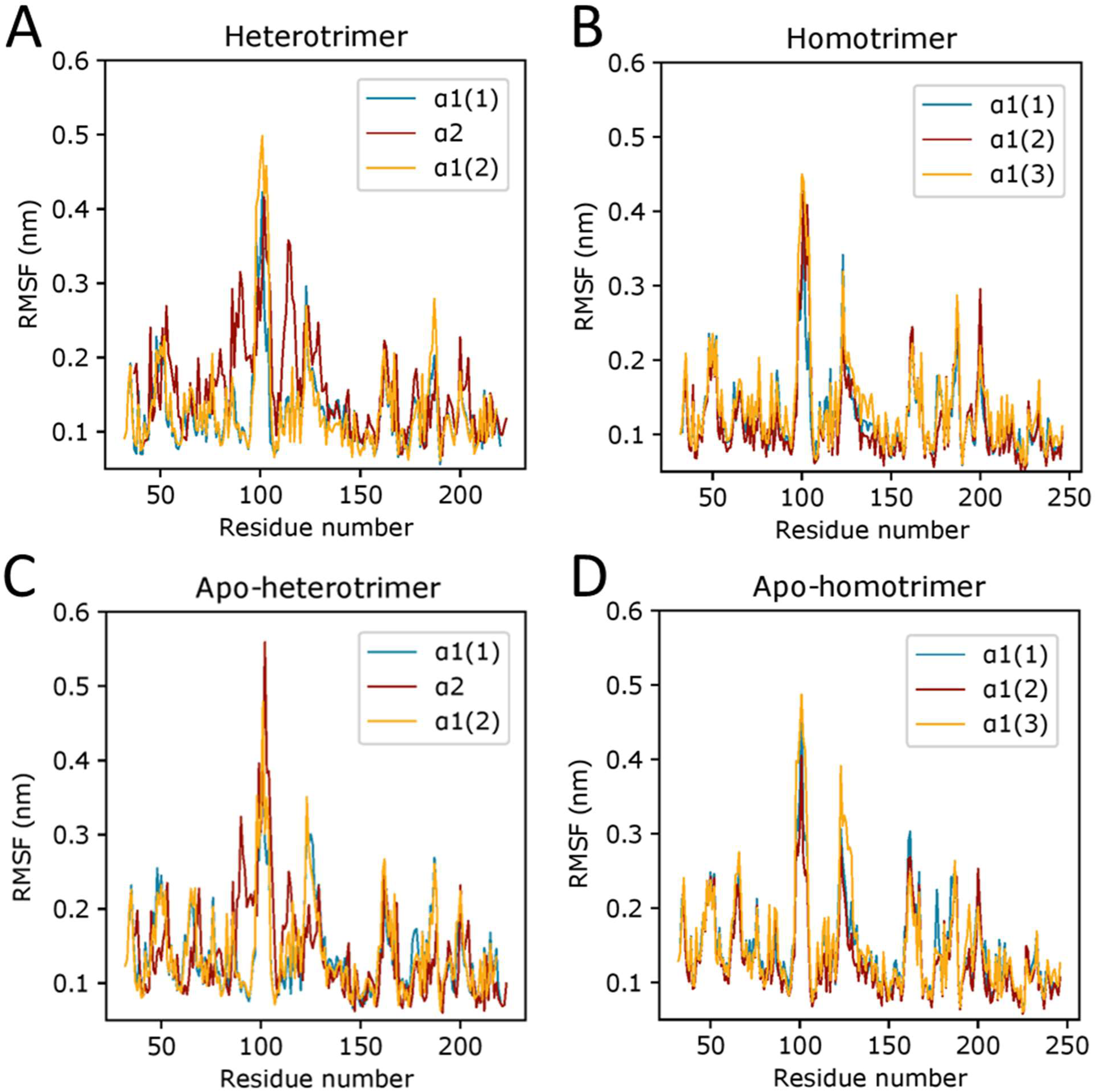
The RMSF of the backbone as a function of amino acids. Each chain is displayed separately per trimer. A: Heterotrimer; B: Homotrimer; C: Apo-heterotrimer, D; Apo-homotrimer.

To capture changes in dynamics that were directly due to calcium loss, RMSF differences between calcium-replete (holo) and calcium deficient (apo) trimers were calculated for both the heterotrimer and homotrimer (Fig. 4). This revealed an overall destabilisation of the alpha-1 chains of the heterotrimer and homotrimer without calcium, indicated by negative values. Conversely the alpha-2 chain of the heterotrimer stabilised without calcium. However, around residue 100 one alpha-1 chain stabilised in both trimers, whilst the alpha-2 chain of the heterotrimer strongly destabilised.

**Figure 4:**
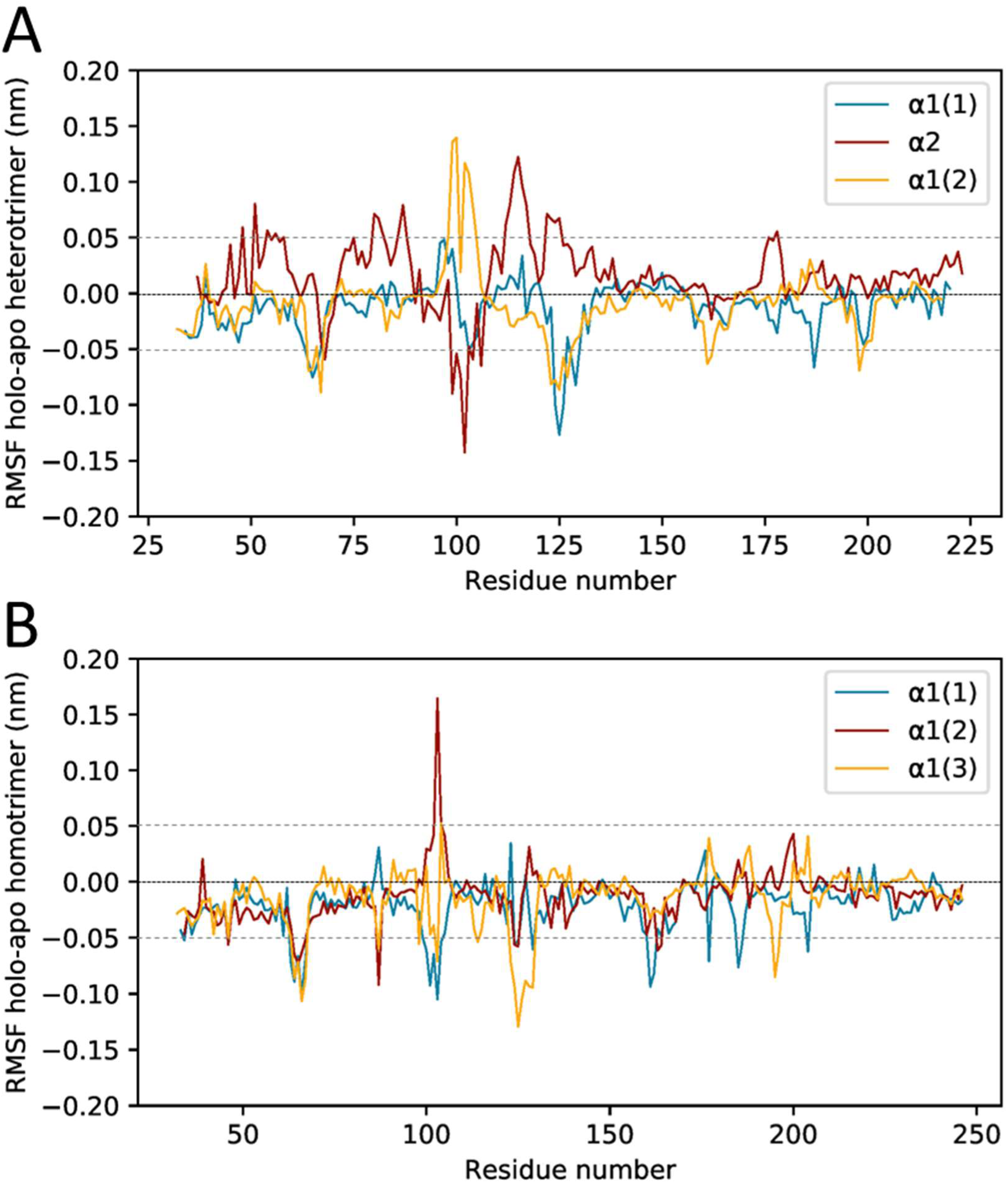
Per-residue RMSF differences between holo- and apo- trimers. The RMSF values for the holo- trimers were subtracted from the apo- trimers and the difference was then plotted per chain for the heterotrimers (A) and homotrimers (B).

To visualise the location of the RMSF hotspots, those residues with RMSF values >0.05 or <-0.05 were highlighted (Figure 5). This confirmed that interface regions in the base of the trimers were among the most perturbed by removal of calcium ions from the structures.

**Figure 5:**
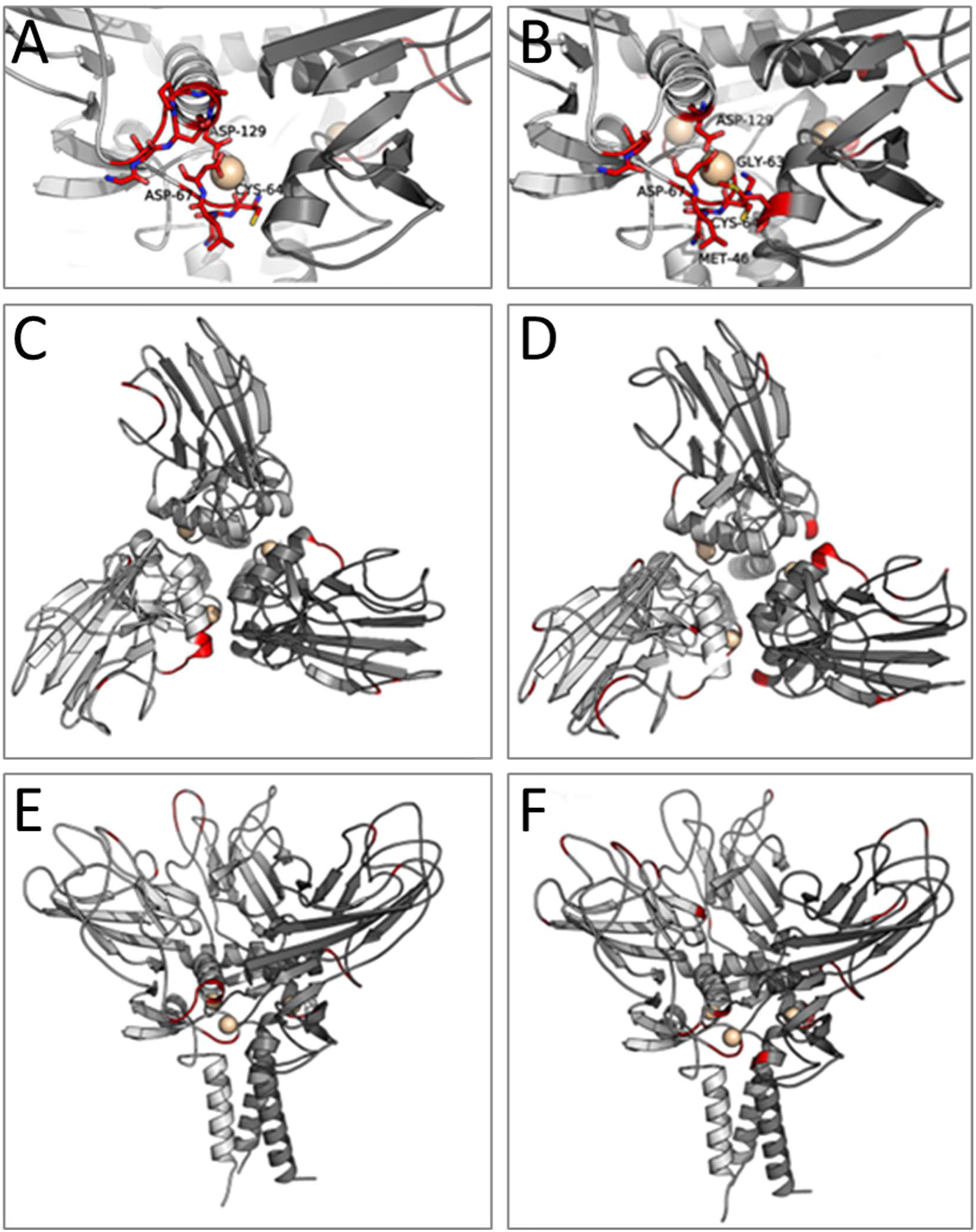
RMSF hotspots in the heterotrimer and homotrimer. A, B: Zoomed view of calcium binding region at the interface of the heterotrimer (A) and homotrimer (B). C, D: Top-down view of the heterotrimer (C) and homotrimer (D). E, F: Side-on view of the heterotrimer (E) and homotrimer (F). The alpha-1(1) chain, then the heterotrimer alpha-2 chain (E) or homotrimer alpha-1(2) (F) and then the alpha-1(3) chain are shaded from lightest to darkest. Hotspots, defined as residues with a difference in RMSF values >0.05 or <- 0.05, are shown in red. Calcium ions are shown as wheat-coloured spheres.

To visualise the impact of calcium loss on the heterotrimer, snapshots of the calcium binding region were taken every 200 ns, starting from 0 ns, for both the holo- (calcium replete) and apo- (calcium depleted) heterotrimers, and then overlaid (Figure 6). All of the intrinsically disordered regions were destabilised, but the calcium-binding loop was most obviously affected. In the holo-protein (Figure 6A), the calcium-binding loop maintained a similar position throughout the trajectory, staying close to the neighbouring chain (Figure 6A). However, in the apo-protein it displayed increased conformational flexibility and moved inward (Figure 6B).

**Figure 6:**
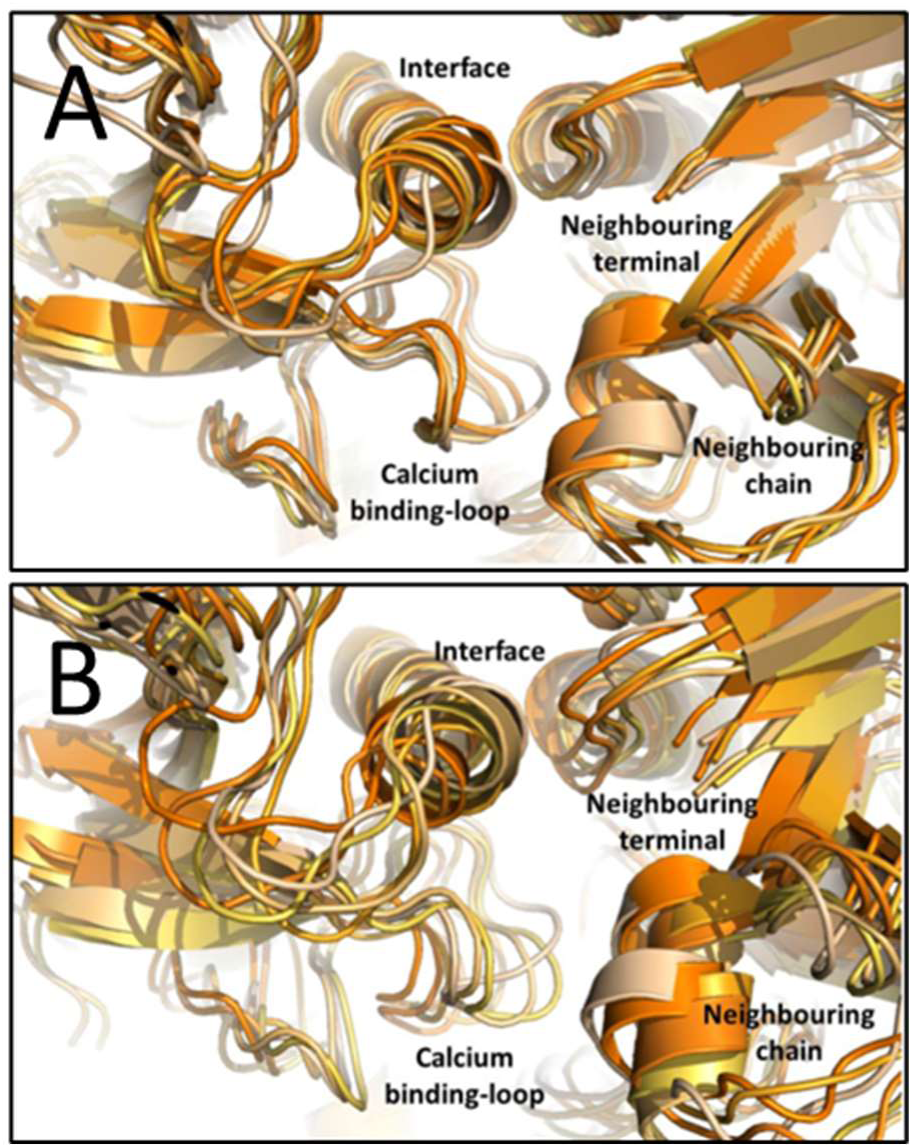
Time evolution of the calcium binding region in heterotrimers. A) Holo-heterotrimer. B) Apo- heterotrimer. Snapshots were taken every 200 ns from 0 to 1000 ns and overlaid. The time points are coloured from the earliest time point in the lightest shade to the latest time point in the darkest shade. The calcium ion is not visualised for the holo-protein for the purpose of visual clarity.

Overall the combination of StructureViz analysis, RMSF values and trajectory visualisation demonstrate a role for structural calcium in co-ordinating hydrogen bond and salt-bridge formation at the interface between chains. Without calcium bound, these interface regions display increased conformational flexibility

### Calcium maintains sufficient proximity for interchain disulphide bonding

It is known that after the individual alpha-chains associate, they become irreversibly disulphide bonded by protein disulphide isomerase (PDI). The cysteine code ensures the proper assembly of homotrimers and heterotrimers and prevents α2(I) homotrimers from forming. In the absence of calcium, no trimers form; only monomers and short-lived dimers are present (18). The relevant cysteines and the α2(I) serine are visualised in Figure 7 A-B. To determine the impact loss of calcium has on covalent bond formation, the distance between the C2 and C3 cysteines, and the α2(I) serine to C3 cysteine for the heterotrimer, at each chain interface was measured (Figure 7 C-F). Pairwise comparisons for linear mixed models fitted to the interchain distances revealed significant differences for the holo-trimer to apo-trimer comparisons for the alpha-1(1):alpha-2 bond distance in the heterotrimer (p=0.018, blue in Figure 7 C &E) and for the homotrimer alpha-1(2):alpha-1(3) and alpha-1(3):alpha-1(1) bone distances (p=0.014 and p<0.001, red and yellow respectively in Figure 7 D & F). For the other comparisons the p-values were not significant (p=0.720, heterotrimer chains B-C Ser--Cys, red; p= 0.116, heterotrimer C-A C2-C3, yellow; p=0.084, homotrimer A-B C2-C3, blue). Disulphide bonds can form between cysteines at distances from 0.3 nm to 0.75 nm (20,21). After the disulphide bond has formed the linkage is typically about 2.04 Å (0.2 nm) in length. At distances greater than 0.8 nm disulphide bonds are unlikely to be observed (20). For the holo-trimers with calcium, the distance between the C2 and C3 cysteines remained near constant through the equilibrium simulations, fluctuating between 0.6 and 0.8 nm (Figure 7 B&C). In contrast for the apo-trimers without calcium (Figure 7 D&E), the C2-C3 the distance increased over the course of the simulation, with average values generally greater than 0.9 nm (Figure 7 E&F). Hence simulations indicate that structural calcium in the trimers maintains a suitable distance between the C2 and C3 cysteines for covalent disulphide bonds to form.

**Figure 7:**
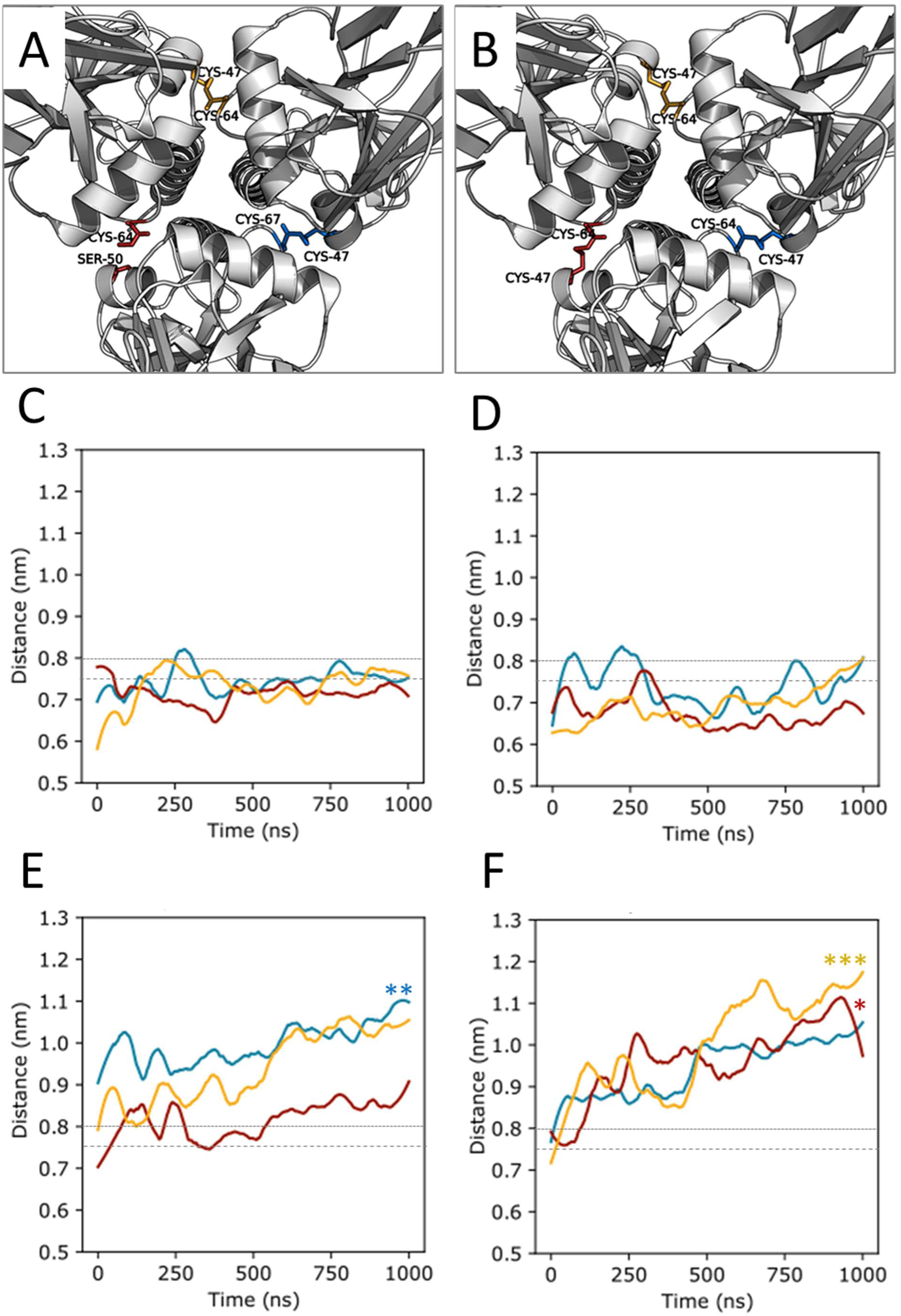
Interchain disulphide bonds and simulated inter-residue distances for the C-propeptide heterotrimer and homotrimer. A, B: Top-down view of the inter-chain disulphide bonding cysteines for the heterotrimer (A) and homotrimer (B). For the heterotrimer (A), the alpha-2:alpha-1(1) chain interface (red) does not contain a disulphide bond, due to the Cys-Ser substitution in the alpha-2(1) chain. The bonds are shown as sticks. The remainder of the chains are shown in light grey as cartoon representations. C-F: Smoothed average distance between inter-chain disulphide bonding residues over the course of the three equilibrium simulation replicates for the heterotrimer (C), the homotrimer (D), the apo-heterotrimer (E) and the apo-homotrimer (F). The alpha-1(1):alpha-2 (heterotrimer) or alpha-1(1):alpha-1(2) (homotrimer) distance is shown in blue, the alpha-2:alpha-1(3) (heterotrimer) or alpha-1(2):alpha-1(3) (homotrimer) distance in red and the alpha-1(3):alpha-1(1) distance in yellow. *P<0.05, **P<0.01 and ***P<0.001 for the holo- to apo-trimer comparisons.

### Calcium is more strongly bound to the alpha-1(I) chain

The calcium ion is co-ordinated by three conserved residues in the calcium binding loop: ASP-59, ASN-61 and ASP-67 in the α1(I) chain, and ASP-62, ASN-64 and ASP-70 in the α2(I) chain. The calcium binding loops have 83% sequence identity and vary by only two residues. ASN-65 and LEU-66 in the α1(I) chain are replaced by THR-68 and MET-69 in the α2(I) chain (Supplementary Figure S1). Differences in calcium binding to each chain were determined using pulling simulations and τ random accelerated molecular dynamics.

### Pulling simulations suggest a higher rupture force but a similar binding affinity, for the calcium ion on the α1(I) than the α2(I) chain

After a short 10 ns equilibrium simulation, run COM pulling simulations were carried out to extract the calcium ion from its binding site (22) (Supplementary Figure S2). The force-time curves for the dissociation of calcium from each chain show an approximately 2-fold difference in the rupture force (F_max_) required to separate the calcium ion from its binding site in the α1(I) chain for the calcium ion compared to the α2(I) chain (Figure 8 A-B). The force profiles generated using Amber ff99SB-ILDN parameters (Fig 6A) appear to demonstrate a three-step unbinding, reflecting the three co-ordinating residues. This is paralleled for the α1(I) chain with partial charges (Fig 6B) though the partial charge profiles are noisier.

**Figure 8:**
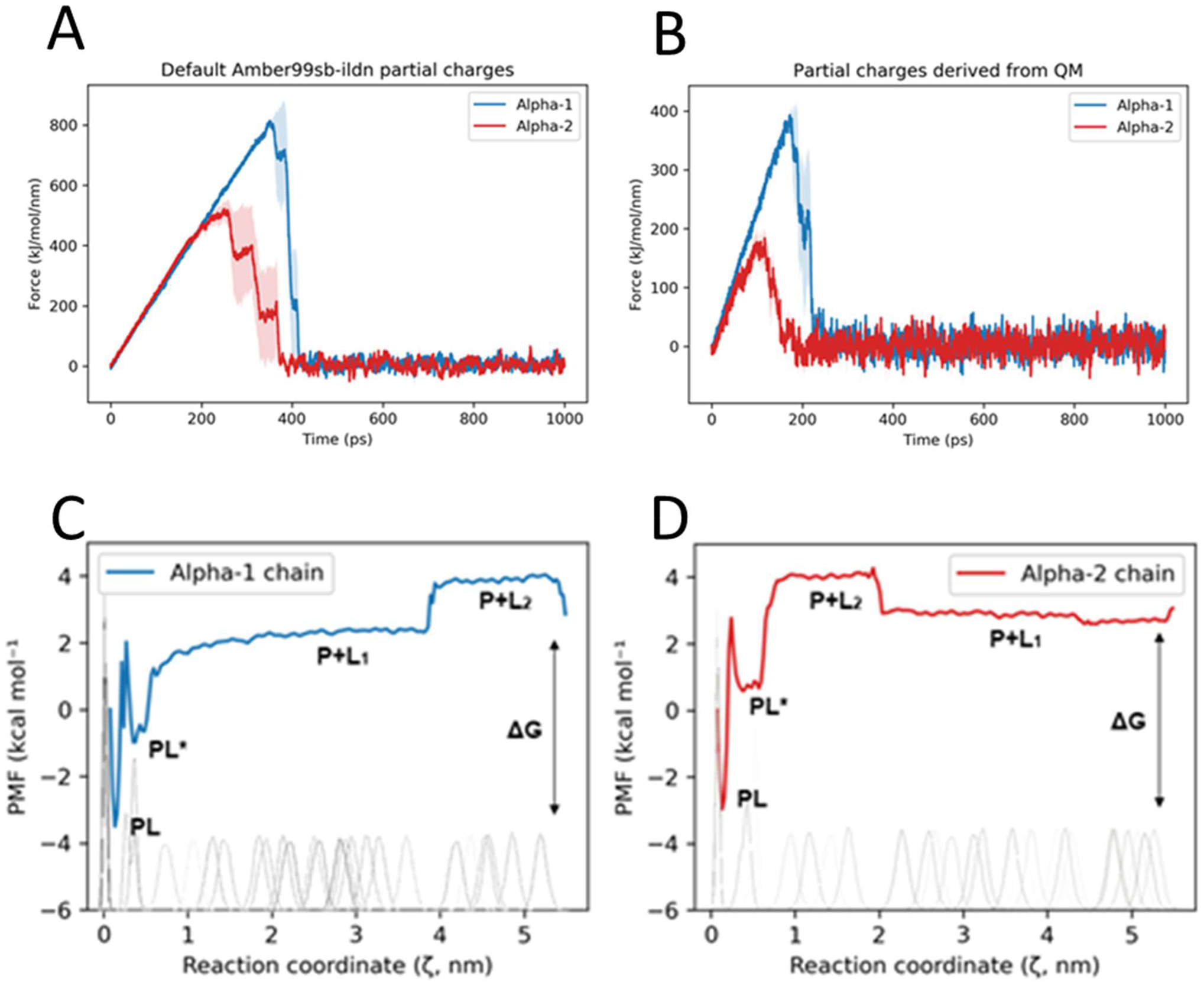
Steered molecular dynamics simulations of the calcium ion uncoupling from its binding site on C-propeptide monomers. A, B: Force-time curves using default Amber ff99SB-ILDN parameters (A), and partial charges determined using QM, calculated by Gaussian 09 (B). The darker lines are the average values over 5 replicates. The shaded areas are the error (standard deviation). C-D: Potential of mean force (PMF) for calcium ion unbinding from the α1(I) chain (C) and the α2(I) chain (D). PL represents the lowest energy ligand binding. P+L(_1-2_) represents the unbound states. PL*represents a possible metastable intermediate bound state, as described in (23). The sampling windows are shown as histograms. The change in Gibbs free energy (ΔG) is also indicated.

Umbrella sampling was then used to compute the binding free energy from the potential of mean force (PMF) for both chains. The unbinding pathway had one potential intermediate step, PL*, before the calcium ion uncoupled from the protein, when the calcium ion then appeared to exist in two states P+L_1_ and P+L_2_ (Fig 8 C-D). The change in Gibbs free energy, ΔG (the binding affinity), the difference between the calcium ion bound to the protein (PL) and the calcium ion in solution P+L_1_, was -5.92 kcal mol⁻¹ for the α1(I) chain and -5.99 kcal mol⁻¹ for the α2(I) chain, with the bound state being more energetically favoured (23). The similar free energies suggest the α1(I) chain and α2(I) chains had similar binding affinity for the calcium ion.

### Structural calcium has a higher residence time in α1(I) chains

τRAMD was employed for cross-validation of the results and to calculate the relative residence times to inform binding kinetics. For each replica a Kolmogorov-Smirnov test was carried to confirm that the measured distribution of dissociation times were within the values reported in the original τRAMD protocol (Table 3 and Supplementary Figure S4). There was a 1.8-fold increase in the residence time (τ) of the calcium ion in the α1(I) chain compared to the α2(I) (Table 4) and a statistically significant difference between the mean relative residence times for each chain (p=0.021).

**Table 3:**
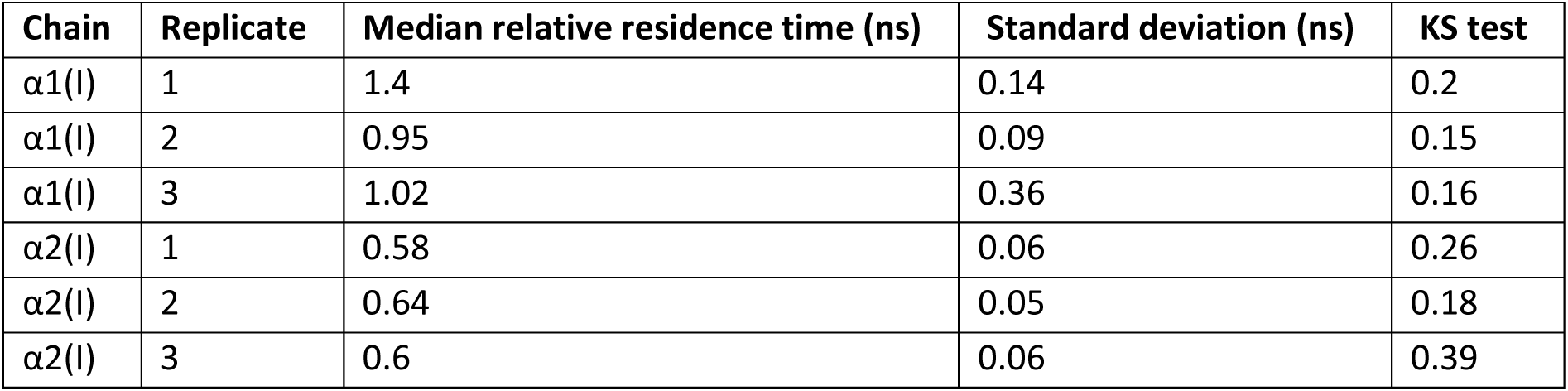
Residence times, relative k_off_ values, and Kolmogorov–Smirnov test results calculated 329 from τRAMD simulations.

| Chain | Replicate | Median relative residence time (ns) | Standard deviation (ns) | KS test |
| --- | --- | --- | --- | --- |
| $\alpha 1(I)$ | 1 | 1.4 | 0.14 | 0.2 |
| $\alpha 1(I)$ | 2 | 0.95 | 0.09 | 0.15 |
| $\alpha 1(I)$ | 3 | 1.02 | 0.36 | 0.16 |
| $\alpha 2(I)$ | 1 | 0.58 | 0.06 | 0.26 |
| $\alpha 2(I)$ | 2 | 0.64 | 0.05 | 0.18 |
| $\alpha 2(I)$ | 3 | 0.6 | 0.06 | 0.39 |

**Table 4:**
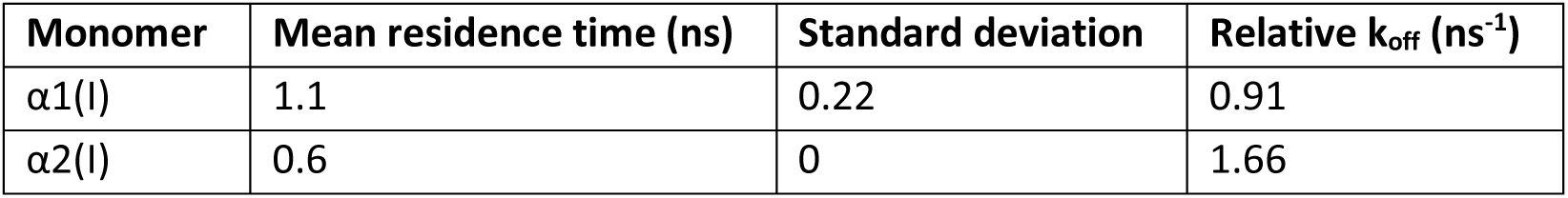
Mean residence times and relative k_off_ values calculated from τRAMD simulations.

| Monomer | Mean residence time (ns) | Standard deviation | Relative $k_{\text{off}}$ ($\text{ns}^{-1}$) |
| --- | --- | --- | --- |
| $\alpha 1(I)$ | 1.1 | 0.22 | 0.91 |
| $\alpha 2(I)$ | 0.6 | 0 | 1.66 |

### Pulling simulations suggest that the α2(I) chain has a higher trimer affinity than a third α1(I) chain in the presence of structural calcium

Steered MD and umbrella sampling was used to determine whether the homotrimeric or heterotrimeric form of the type I collagen C-propeptide, including a single alpha-2(I) chain is energetically favoured. The force-time curves for an α2(I) or corresponding α1(I) chain separating from a heterotrimer and homotrimer at 0 ns and after 1000 ns of MD were compared. At 0 ns, the results suggested the holo-heterotrimer had the strongest binding affinity, followed by the holo-homotrimer, then the apo-trimers (Figure 9A). The heterotrimer appeared to display a two-step unbinding process, with a smaller rupture event after the first large rupture event. This was reflected in the number of hydrogen bonds between the holo-heterotrimer; after an initial sharp drop, the number of bonds plateaued until the 500 ps mark (Figure 9B). After 1000 ns of equilibrium MD the same experiment was carried out. However, this time, the holo-heterotrimer and apo-heterotrimer displayed the highest binding affinities, followed by the holo-homotrimer and holo- heterotrimer (Figure 9C).

**Figure 9:**
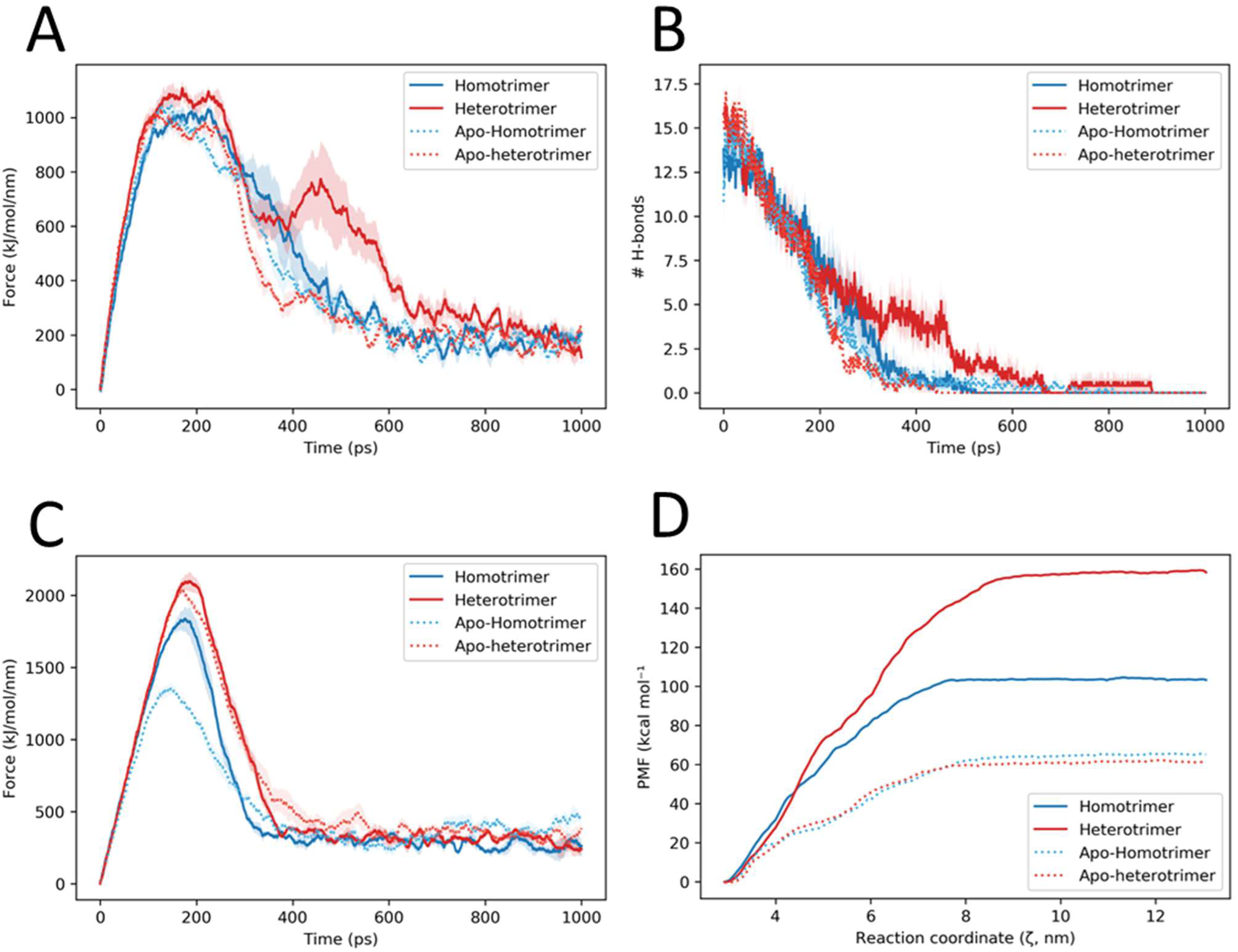
Steered molecular dynamics simulations of the α2(I) or corresponding α1(I) chain pulled from C- propeptide heterotrimers or homotrimers. A: Force-time curves of type I collagen trimers with and without calcium bound (holo- and apo-) after 0 ns of equilibrium and B: the corresponding hydrogen bond plot. C: Force-time curves of type I collagen trimers pulled after 1000 ns of equilibration. D: Potential of mean force (PMF) curves for, heterotrimer, apo-heterotrimer homotrimer and apo-homotrimer.

The 0 ns time point pulling simulations were selected for umbrella sampling. The histograms initially demonstrated that additional windows were required at the early time points, but all systems were then sufficiently sampled (Supplementary Figure S5). The umbrella sampling results suggest that the holo- heterotrimer has a higher binding affinity than the holo-homotrimer and that both isoforms have a much higher binding affinity than their corresponding apo- versions (Figure 9D and Table 5). The apo- homotrimers and heterotrimers displayed little-no difference in their binding affinities, in keeping with them both being equally unfavourable when there is a complete lack of calcium

**Table 5:** Binding affinities of apo- and holo- type I collagen trimer isoforms.

| Trimer | Binding affinity ($\Delta G$) |
| --- | --- |
| Homotrimer | 104.52 |
| Heterotrimer | 159.40 |
| Apo-homotrimer | 66.13 |
| Apo-heterotrimer | 62.70 |

## Discussion

In the present study we have uncovered a calcium-dependent mechanism that determines whether type I collagen forms heterotrimeric or homotrimeric. The findings of the simulations show that loss of calcium affects the hydrogen bonding network at the chain interfaces, meaning that partner cysteines are too far apart to form disulphide bonds. The structural calcium ion is more strongly bound to the alpha-1(I) chain than the alpha-2(I) chain, meaning that even though the heterotrimer is energetically favoured, it would be less able to form if free calcium ions were sequestered by alpha-1(I) chains. We propose a model whereby reduced a reduced calcium ion concentration in the endoplasmic reticulum would favour homotrimerisation over heterotrimerisation of type I procollagen (Figure 10).

**Figure 10.**
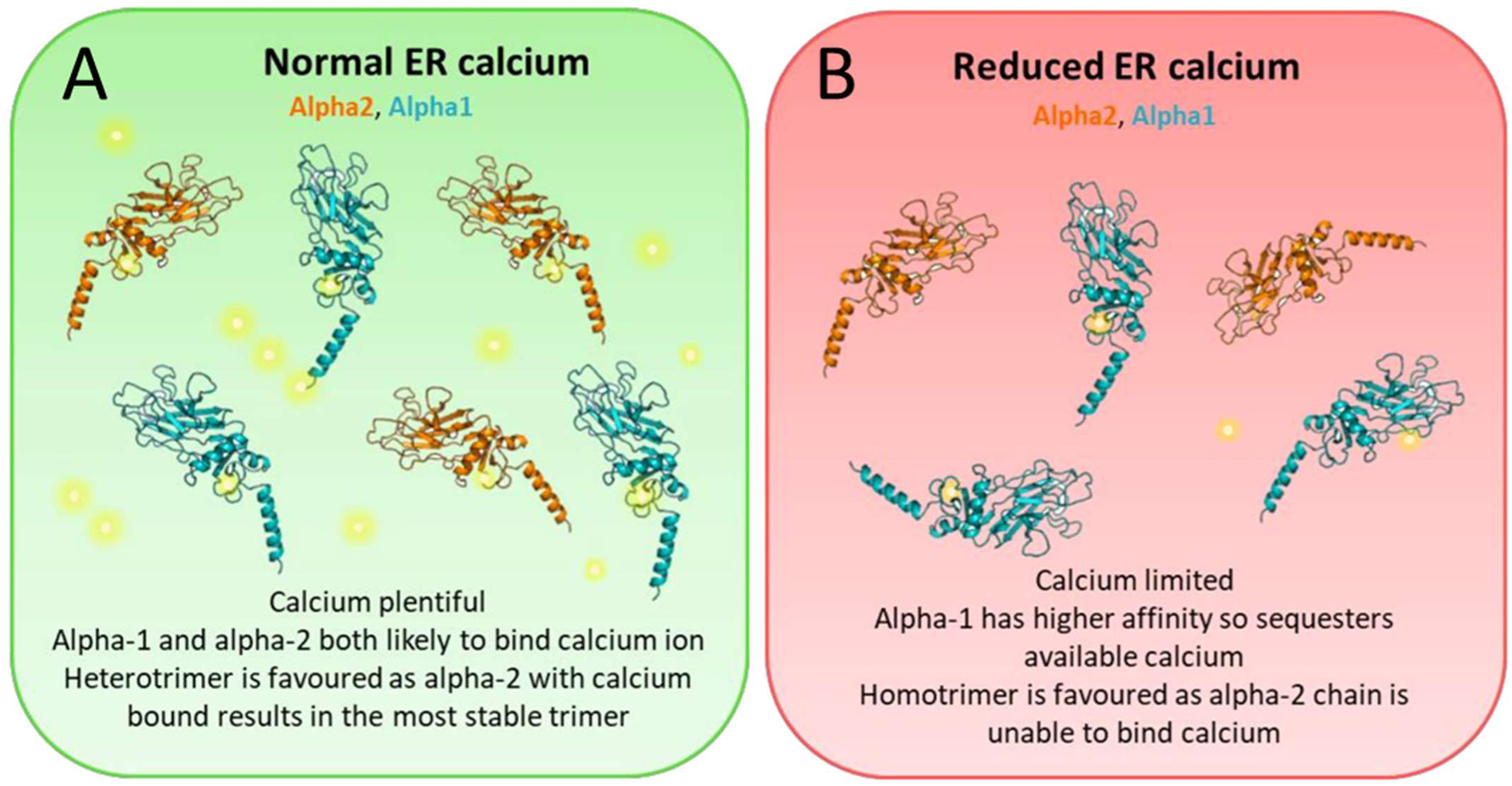
Model for calcium regulation of type I collagen hetero-versus homotrimerisation. A: Heterotrimers are favoured in normal ER calcium concentrations, where there is sufficient calcium to bind all pro-alpha-1(I) and pro-alpha-2(I) monomers. B: Homotrimers are favoured when calcium concentrations are limited as the requisite calcium ions are mopped up by alpha-1(I) chains, leaving alpha-2(I) chains unable to join trimers. For clarity, only the C-propeptides of the procollagen alpha-1(I) and alpha-2(I) chains are shown; triple helical regions and N-propeptides are omitted. Other ER chaperones required for procollagen folding are not shown. The alpha-1(I) C-propeptide monomers are shown in blue and the alpha- 2(I) monomers in orange. Calcium ions are depicted as yellow luminous spheres.

Previous studies have shown that the homotrimer will form in the absence of alpha-2(I) chains due to genetic inactivation or epigenetic silencing of COL1A2 (12,24,25), or by over-production of the alpha-1(I) chain, either artificially or due to a common COL1A1 polymorphism (26,27). However gene dosage and the relative abundances of COL1A1 and COL1A2 mRNAs, do not directly predict heterotrimerisation versus homotrimerisation (28,29), This is likely because COL1 mRNA translation is coordinated by numerous cytosolic factors (30) and the alpha-2(I) chain sequence itself favours heterotrimerisation.

Collagens are co-translationally translocated into the lumen of the rough endoplasmic reticulum (rER) during biosynthesis. The entire chain must be translated before the C-propeptide can fold and mediate trimersation. Normal free ER calcium concentrations range from 0.5 - 2.0 mM in many different cell types. Calcium pumps and channels control the uptake or release of calcium from the ER to the cytoplasm and store-operated entry maintains equilibrium calcium levels, ensuring that cytosolic signalling does not compromise ER function (31). The amount of calcium available in the ER is buffered by calcium-binding proteins, primarily calsequestrin in skeletal and cardiac muscle, and calrecticulin in other tissues. Calcium-dependent collagen chaperones such as BiP/GRP78, GRP94, protein disulphide isomerase (PDI) and calnexin also act as calcium stores (32). The agonist-releasable calcium concentration in the ER estimated to be approximately 1 mM (33). Previous sedimentation-equilibration experiments with recombinant C- propeptides lacking cysteines, showed that homotrimerisation occurred in 0.5 mM calcium, but that monomers predominated in the absence of calcium (18). Future studies with mixtures of alpha-1(I) and alpha-2(I) chains over a range of calcium concentrations could elucidate the minimum calcium concentration at which heterotrimerisation occurs.

Chronic decreases in ER calcium concentrations can affect the unfolded protein response (UPR) leading to increases in misfolded proteins and subsequent apoptosis (32). The importance of intracellular calcium homeostasis is demonstrated by the ER stress and dysregulated type I collagen synthesis caused by loss of TMEM38B, an ER membrane monovalent cation channel, which results in osteogenesis imperfecta (OI) (34). However, ER calcium concentrations can be decreased in many pathological states including diabetes, ischemia, cardiovascular disease, viral infections, asthma, liver disease and cancer (32). Calcium homeostasis also becomes dysregulated in ageing due to oxidative damage, decreased expression of the ER calcium ion pump SERCA and alterations in calcium-sensing proteins (35-37). Hence age- and disease-associated decreases in ER calcium could lead to formation of type I collagen homotrimer, rather than heterotrimer. Chronic diseases can be associated with increased focal collagen production, and the corresponding production of the homotrimeric form could accelerate fibrosis, due to the increased resistance of the homotrimer to MMP-mediated turnover.

Fibroblasts characteristically produce abundant type I collagen, particularly during development and tissue homeostasis. In fibroblasts, calcium signalling is mechano-sensitive and responds to the cell’s 3D environment (38), which is especially relevant for proliferation and migration of fibroblasts during wound healing and leads to cyclic fluctuations in the amount of free calcium available (39). Hence the chain-specific trimerisation of type I collagen may be influenced by the mechanical environment.

In two diseases of the skin, Darier disease and Hailey-Hailey disease, the ER calcium ion pumps SERCA2 and SPCA1 become mutated, respectively, leading to dysregulation of calcium homeostasis and phenotypic features such as the formation of warts, lesions, blisters and malodorous plaques (40). It is unknown if homotrimeric type I collagen is present in these conditions, though a COL1A1-PDGFB fusion gene is implicated in similar skin conditions (41).

Type I collagen homotrimers promote metastasis and invasion in cancer cells by providing MMP resistant pathways for cell migration (6,25). ER calcium is often decreased in cancer cells, which can create resistance to apoptosis (42). It may therefore be that this depleted ER calcium contributes to homotrimerisation of type I collagen during malignant transformation.

Considering the hydrogen bonding network of the C-propeptide trimers, it was noted that the homotrimer lacked an inter-chain hydrogen bond between ASP-43.B (on the alpha-1(2) chain) and CYS-64.C (on the alpha-1(3) chain) throughout two independent simulations, even though a C2-C3 disulphide bond still forms at that position and all three chains have the same structure. This asymmetry in the homotrimer is not entirely unprecedented, as the type III collagen C-propeptide displays intrinsic asymmetry despite also being a homotrimer (16). Despite not forming a C2-C3 bond at this interface the heterotrimer did have a hydrogen bond between the homologous ASP-46.B (on the alpha-1(2) chain) and CYS-64.C (on the alpha- 1(3) chain); though the bond between ASP-43.C (on the alpha-1(3) chain) and CYS-64.A (on the alpha-1(1)) chain was weaker. The results from the homotrimer and heterotrimer suggest that inter-chain ASP-43 and CYS-64 hydrogen bonding is not necessary at every chain interface for trimerisation to occur, and that the heterotrimer may have hydrogen bonds that compensate for the lack of C2 residue in the α2(I) chain.

In addition to providing general structural insight, uncovering the hydrogen bonding network demonstrates how some known type I collagen C-propeptide mutations could cause disease. Around 6.5% of OI patients have mutations in the C-propeptide region. The OI causing mutation P-1182-R (at position 63 in the α2(I) C-propeptide) is proximal to the ASN-64 residue (43,44). This residue is important in stabilising the binding interface between chains, so replacing the neighbouring small and neutrally charged proline with a bulky, hydrophilic, positively charged arginine will likely interfere with hydrogen bond formation and chain incorporation. The same principle applies for other OI causing mutations. THR-1431 (located at position 213 in the α1(I) C-propeptide) is close to the ARG-218 residue that participates in hydrogen bonding with GLU-177 in the neighbouring chain. TYR-1263 (located at position 144 in the α2(I) C-propeptide) (43) has two neighbouring residues that participate in inter- and intra-chain hydrogen bonding respectively. It may also be that these mutations interfere with the solvent accessible area or directly repel their neighbours.

Overall the combination of RMSF values, trajectory visualisation, and StructureViz analysis demonstrated a role for structural calcium in co-ordinating hydrogen bond and salt-bridge formation at the interface. Without calcium bound, these interface regions display increased conformational flexibility and could be less likely to form the bonds necessary for trimerisation *in vivo*. It seems the structural calcium typically serves to guide the C2 and C3 cysteines towards each other and without it covalent bonding is unlikely to take place. This likely explains why there are only short-lived dimer species observed in expression systems that lack calcium in solution. One limitation of the present study is that it is solely computational, whilst sedimentation-equilibration experiments with mutated recombinant C-propeptides could validate the findings *in vitro*. A further limitation is that the structure of the heterotrimer is based on homology modelling, rather than a crystal structure, which is because the heterotrimeric C-propeptide could not be crystallised (5). However homology modelling is more reliable than other methods could be employed here due to the high sequence homology between the alpha-chains. It may be that future advances in protein crystallization techniques or protein modelling may allow the model to be refined and retested. Finally the simulations pulling a monomer from a C-propeptide trimer could only be carried out once, due to the computationally intensive nature of umbrella sampling. Future advances in high performance computing and GPU capabilities could allow this approach to be repeated. Such advances may also permit modelling of the folding of the entire procollagen triple helix and to define how specific amino acid substitutions affect protein folding rate and the final folded structure.

## Experimental procedures

### Generating homology models

The previously published structure of the C-propeptide of homotrimeric type I collagen was used as a template for homology modelling (PDB accession number: 5K31, biological assembly 3) (5). Two residues had been mutated in the original structure for crystalisation purposes; these were mutated to match the canonical sequence using PyMOL’s mutagenesis feature and incomplete sidechains were built using SWISS-PDB viewer. Glycerol and an excess chloride ion were also stripped. SWISS-MODEL (45), was used to create the homology model for the C-propeptide of heterotrimeric type I collagen using the modified homotrimeric C-propeptide structure as a template. To create apo- versions of the proteins, the remaining structural ions were also stripped. All trimers were simulated with the interchain disulphide bonds reduced. Monomers were created by extracting an α1(I) chain or the corresponding α2(I) chain.

### Force fields and conventional simulation protocol

GROMACS 2019.3 (46,47) was used to run all simulations of type I collagen trimers and monomers in solution. The AMBER99SB-ILDN force field (48) was used to describe the system topology. Three replicates were carried out. Hydrogens were replaced with virtual sites to allow for a longer time step of 5 fs, and sampling of longer timescales (49). A fourth replicate with explicit hydrogens was also carried out.

Standard simulation set-ups and protocols associated with the AMBER99SB-ILDN force field were used to carry out the MD calculations: electrostatic interactions were calculated using the PME (particle-mesh Ewald) method and short-range non-bonded interactions were cut off at 1.0 nm (50,51). Verlet neighbour search was used with a screening constant of 20 fs (52). The LINCS algorithm was used to constrain all bonds during the simulations (53).

The TIP3P water model was used to solvate the system. The systems were then neutralised using potassium ions. Additional potassium and chloride ions were then added to simulate a physiological salt concentration of 150 mM. Potassium was selected as the cation to best represent the conditions inside the endoplasmic reticulum (ER). Energy minimisation was carried out using the steepest descent algorithm and terminated after 50,000 steps or when the maximum force was reduced to 1000 kJ mol^−1^ nm^−2^. Equilibration was carried out in two steps; the first step was a constant volume (NVT) ensemble where the protein and aqueous phase (water plus ions) were coupled to separate temperature baths at 310 K using the modified Berendsen thermostat (V-rescale). The second step was a constant pressure (NPT) equilibration with the Parrinello-Rahman barostat to maintain the pressure isotropically at 1.0 bar. Production MD was then carried out for each replicate under the same NPT ensemble for 1000 ns (resulting in 16,000 ns of trajectory data in total).

### Enhanced sampling protocols

Enhanced sampling methods were applied to type I collagen C-propeptide monomers and trimers. For trimers, protein and structural ion (where appropriate) co-ordinates were extracted at the beginning and end of the equilibrium simulations. To investigate the affinity of an α1(I) or α2(I) chain for its neighbouring two chains τRAMD, steered MD and umbrella sampling were employed. τRAMD was used to optimise sampling of the collective variable, by pulling a mobile chain in a random direction from two stationary chains. The τRAMD standard protocol was run 25 times for each replicate. After sampling the unbinding pathway, steered MD (in this case, centre-of-mass (COM) pulling) was used to pull the mobile chain away from the two stationary chains for all the trimers of interest. The trimers were placed in a large rectangular box, solvated, neutralised, energy minimised and equilibrated following the same protocol as the equilibrium simulations. The trimer was rotated so the mobile chain was orientated in the direction of the unbinding pathway. Backbone restraints were applied to the two stationary chains. Pulling specific regions of the protein confirmed the importance of the two interface regions for stabilising the bonds between the chains (Supplementary Figure S7). However, to conserve all the information about the unbinding pathway, the pull force was applied across the whole protein. A spring constant of 1,500 mol^−1^ nm^-2^ was applied with a pull rate was 0.05 nm ps^−1^ (Supplementary Figure S7). For each trimer replicate, ten COM pulling simulations were carried out. An average force vs time plot was calculated for each snapshot, along with the average rupture force. Sample pulling trajectories were then used to generate starting configurations for umbrella sampling. Snapshots were taken every 0.1-0.2 nm along the collective variable up to a final centre-of-mass distance of approximately 12 nm, resulting in roughly 50 windows for each replicate. 10 ns of MD were carried out per window to ensure sufficient sampling. The results were then analysed using the weighted histogram method (WHAM).

To investigate the binding affinity of the structural calcium for a single α1(I) or α2(I) chain, the uncoupling of the calcium ion from the binding loop was studied, employing the same techniques. A shorter production MD run (10 ns) was carried out for α1(I) and α2(I) monomers (three replicates) and these end- state co-ordinates used for τRAMD and steered MD/umbrella sampling. For each replicate 25 τRAMD runs were carried out. A pulling force of 500 kJ mol^−1^ nm^−2^ was applied to the ion in a random direction. The simulation terminated after the ion moved 5 nm from the binding site. The relative residence time (τ_comp_) was then calculated as the simulation time it takes for the ligand to have dissociated in 50% of the trajectories. For the steered MD of monomers, the calcium-binding loop of the α1(I) and α2(I) chains served as the reference group (Supplementary Figure S2 B&C). The backbone atoms of the loop were position restrained, allowing the side chains to move freely. The calcium ion was pulled away from the loop along the z-axis using a spring constant of 500 kJ mol^−1^ nm^−2^ and a pull rate of 0.005 nm ps^−1^. Snapshots were taken every 0.2 nm along the collective variable up to a final centre-of-mass distance of approximately 5 nm, resulting in approximately 30 windows for each replicate. As with the trimers 10 ns of MD was carried out per window and analysed using the WHAM method. We also repeated these simulations with partial charges derived via quantum mechanics using the DFT-B3LYP hybrid functional with the 6-31G(d,p) basis set to negate any forcefield effects (Supplementary Figure S2D).

### Analysis

For the equilibrium simulations proteins were made whole and jumps and PBC were removed, then, translational and rotational movements were also removed. Trajectories were visualised in VMD, and single frames were for visualised using PyMOL (Version 2.0 Schrödinger, LLC). The inbuilt *gmx rmsf*, *gmx gyrate*, *gmx distance* and *gmx rmsd* modules were used to analyse the results of the simulations.

Statistical analysis of these outputs was carried out using the R statistical programming environment. Linear mixed models were fit to the data to model the evolution of each trimer over time (using a trimer*time interaction term). A random intercept term was used to estimate the variability between replicates in the study and account for repeated measures. Pairwise comparisons between the trimers were carried out using the *emmeans* package with the Kenward-Roger method. The results of the equilibrium simulations and COM-pulling/steered MD analysis were visualised using the python package Matplotlib. The τRAMD results were analysed using a freely available script (https://github.com/DKokh/tauRAMD).

### Hardware

All simulations were performed on the Barkla high-performance computing cluster located at the University of Liverpool. 10 nodes were made available for the calculations, each with 40 cores and a total of 384 GB of memory. Visualisation and analysis were carried out on a local machine with a 64-bit operating system and an Intel® Core™ i5-3570K processor. 8 GB RAM was available for calculations and graphical manipulations.

## Data availability

The data can be obtained from Emily J Johnson (Emily.Johnson@liverpool) or from the corresponding author on request.

## Supporting information

This article contains supporting information.

## Author contributions

Conceptualization: E.J.J., E.G.C.-L.; Data curation: E.J.J. Formal Analysis: E.J.J., J.V.d.S, Funding acquisition: E.G.C.-L.; Investigation: E.J.J., J.V.d.S, Project administration: E.G.C.-L.; Supervision: A.K.B., E.G.C.-L.; Visualization: E.J.J., Writing – original draft: E.J.J., E.G.C.-L.; Writing – review & editing: E.J.J., J.V.d.S, A.K.B., E.G.C.-L

## Funding and additional information

This work was funded by the Biotechnology and Biological Sciences Research Council (BBSRC), UK [BB/M011186/1, 1945098]

---

**Supplementary Table 1.** Notable hydrogen bonds in the homotrimer and heterotrimer. Alpha-1(1) is denoted chain A, alpha-2 in the heterotrimer and alpha-1(2) in the homotrimer are denoted chain B, whilst alpha-1(3) is denoted chain C. The weight corresponds to how conserved the bond was throughout the simulation, with higher weights corresponding to the most conserved bonds and lower weights corresponding to more transient bonds.

| Trimer | Bond | Position in full length chain | Inter-chain/<br>intra-chain | Weight |
| --- | --- | --- | --- | --- |
| Homotrimer | ARG 42.A - ASP 129.B | ARG 1260.A - ASP 1347.B | Inter-chain | 2.00794 |
| Homotrimer | ARG 39.B - ASN 61.C | ARG 1257.B - ASN 1279.C | Inter-chain | 1.48413 |
| Homotrimer | ARG 42.B - ASP 129.C | ARG 1260.B - ASP 1347.C | Inter-chain | 1.1746 |
| Homotrimer | ASP 43.A - CYS 64.B | ASP 1261.A - CYS 1282.B | Inter-chain | 1.09524 |
| Homotrimer | ALA 128.C - ARG 42.B | ALA 1346.C - ARG 1260.B | Inter-chain | 0.650794 |
| Homotrimer | ARG 42.C - ASP 129.A | ARG 1260.C - ASP 1347.A | Inter-chain | 0.579365 |
| Homotrimer | ALA 128.A - ARG 42.C | ALA 1346.A - ARG 1260.C | Inter-chain | 0.515873 |
| Homotrimer | ARG 42.A - ASP 67.B | ARG 1260.A - ASP 1285.B | Inter-chain | 0.460317 |
| Homotrimer | ASP 43.C - CYS 64.A | ASP 1261.C - CYS 1282.A | Inter-chain | 0.444444 |
| Homotrimer | ARG 39.C - ASN 61.A | ARG 1257.C - ASN 1279.A | Inter-chain | 0.380952 |
| Homotrimer | ARG 39.A - ASN 61.B | ARG 1257.A - ASN 1279.B | Inter-chain | 0.301587 |
| Homotrimer | ARG 39.B - GLN 62.C | ARG 1257.B - GLN 1280.C | Inter-chain | 0.301587 |
| Homotrimer | CYS 64.C - MET 46.B | CYS 1282.C - MET 1264.B | Inter-chain | 0.246032 |
| Homotrimer | PHE 245.A - THR 80.A | PHE 1463.A - THR 1298.A | Intra-chain | 2.01587 |
| Homotrimer | ILE 58.A - ILE 69.A | ILE 1276.A - ILE 1287.A | Intra-chain | 2 |
| Homotrimer | TYR 56.C - VAL 71.C | TYR 1274.C - VAL 1289.C | Intra-chain | 2 |
| Homotrimer | ILE 58.B - ILE 69.B | ILE 1276.B - ILE 1287.B | Intra-chain | 2 |
| Homotrimer | ILE 58.C - ILE 69.C | ILE 1276.B - ILE 1287.B | Intra-chain | 2 |
| Homotrimer | TYR 56.A - VAL 71.A | TYR 1274.A - VAL 1289.A | Intra-chain | 2 |
| Homotrimer | TYR 56.B - VAL 71.B | TYR 1274.B - VAL 1289.B | Intra-chain | 1.98413 |
| Homotrimer | CYS 41.B - THR 80.B | CYS 1259.B - THR 1298.B | Intra-chain | 1.7619 |
| Homotrimer | CYS 41.A - THR 80.A | CYS 1259.A - THR 1298.A | Intra-chain | 1.68254 |
| Homotrimer | CYS 41.C - THR 80.C | CYS 1259.C - THR 1298.C | Intra-chain | 1.61905 |
| Homotrimer | ARG 39.B - ASP 43.B | ARG 1257.B - ASP 1261.B | Intra-chain | 1.53968 |
| Homotrimer | ASP 43.B - CYS 47.B | ASP 1261.B - CYS 1265.B | Intra-chain | 1.38889 |
| Homotrimer | ASN 61.B - GLN 133.B | ASN 1279.B - GLN 1351.B | Intra-chain | 1.24603 |
| Homotrimer | ASP 43.C - THR 40.C | ASP 1261.C - THR 1258.C | Intra-chain | 1.23016 |
| Homotrimer | ASP 43.B - THR 40.B | ASP 1261.B - THR 1258.B | Intra-chain | 1.14286 |
| Homotrimer | ASN 61.A - GLN 133.A | ASN 1279.A - GLN 1351.A | Intra-chain | 1.05556 |
| Homotrimer | ASN 61.C - GLN 133.C | ASN 1279.C - GLN 1351.C | Intra-chain | 1.03968 |
| Homotrimer | ASP 43.C - CYS 47.C | ASP 1261.C - CYS 1265.C | Intra-chain | 0.968254 |
| Homotrimer | ASP 43.A - THR 40.A | ASP 1261.A - THR 1258.A | Intra-chain | 0.912698 |
| Homotrimer | ASN 61.B - ASP 67.B | ASN 1279.B - ASP 1285.B | Intra-chain | 0.888889 |
| Homotrimer | ARG 42.B - LEU 246.B | ARG 1260.B - LEU 1464.B | Intra-chain | 0.873016 |
| Homotrimer | ARG 39.C - PRO 60.C | ARG 1257.C - PRO 1278.C | Intra-chain | 0.865079 |
| Homotrimer | ASN 61.A - ILE 132.A | ASN 1279.A - ILE 1350.A | Intra-chain | 0.857143 |
| Homotrimer | ASN 61.C - ASP 67.C | ASN 1279.C - ASP 1255.C | Intra-chain | 0.84127 |
| Homotrimer | ASN 61.A - ASP 67.A | ASN 1279.A - ASP 1255.A | Intra-chain | 0.84127 |
| Homotrimer | ARG 42.C - MET 46.C | ARG 1260.C - MET 1264.C | Intra-chain | 0.785714 |
| Homotrimer | ARG 42.B - THR 142.B | ARG 1260.B - THR 1360.B | Intra-chain | 0.769841 |
| Homotrimer | ASP 43.A - CYS 47.A | ASP 1261.A - CYS 1265.A | Intra-chain | 0.746032 |
| Homotrimer | ARG 42.A - MET 46.A | ARG 1260.A - MET 1264.A | Intra-chain | 0.68254 |
| Homotrimer | ARG 42.C - LEU 246.C | ARG 1260.C - LEU 1464.C | Intra-chain | 0.674603 |
| Homotrimer | ARG 39.C - ASP 43.C | ARG 1257.C - ASP 1261.C | Intra-chain | 0.650794 |
| Homotrimer | ARG 39.A - ASP 43.A | ARG 1257.A - ASP 1261.A | Intra-chain | 0.619048 |
| Homotrimer | ARG 42.B - MET 46.B | ARG 1260.B - MET 1264.B | Intra-chain | 0.595238 |
| Homotrimer | ARG 42.C - THR 142.C | ARG 1260.C - THR 1360.C | Intra-chain | 0.5 |
| Homotrimer | ARG 39.B - GLN 62.B | ARG 1257.B - GLN 1280.B | Intra-chain | 0.468254 |
| Homotrimer | ASN 61.C - ASP 59.C | ASN 1279.C - ASP 1277.C | Intra-chain | 0.468254 |
| Homotrimer | ARG 39.B - PRO 60.B | ARG 1257.B - PRO 1278.B | Intra-chain | 0.34127 |
| Homotrimer | ASN 61.B - ILE 132.B | ASN 1279.B - ILE 1350.B | Intra-chain | 0.293651 |
| Heterotrimer | ARG 42.A - GLU 130.B | ARG 1260.A - GLU 1249.B | Inter-chain | 2.45238 |
| Heterotrimer | ARG 39.A - ASN 64.B | ARG 1257.A - ASN 1183.B | Inter-chain | 1.59524 |
| Heterotrimer | ASP 43.A - CYS 67.B | ASP 1261.A - CYS 1186.B | Inter-chain | 1.26984 |
| Heterotrimer | ARG 42.C - ASP 129.A | ARG 1260.C - ASP 1347.A | Inter-chain | 0.984127 |
| Heterotrimer | ARG 39.C - ASN 61.A | ARG 1257.C - ASN 1279.A | Inter-chain | 0.984127 |
| Heterotrimer | ARG 42.B - ASN 61.C | ARG 1161.B - ASN 1279.C | Inter-chain | 0.896825 |
| Heterotrimer | ASP 46.B - CYS 64.C | ASP 1165.B - CYS 1282.C | Inter-chain | 0.746032 |
| Heterotrimer | ARG 45.B - ASP 129.C | ARG 1164.B - ASP 1347.C | Inter-chain | 0.587302 |
| Heterotrimer | ARG 45.B - ASP 67.C | ARG 1164.B - ASP 1285.C | Inter-chain | 0.103175 |
| Heterotrimer | ASP 43.C - CYS 64.A | ASP 1261.C - CYS 1282.A | Inter-chain | 0.103175 |
| Heterotrimer | ASP 43.A - THR 40.A | ASP 1261.A - THR 1258.A | Intra-chain | 2.15873 |
| Heterotrimer | ARG 42.A - LEU 246.A | ARG 1260.A - LEU 1464.A | Intra-chain | 2.01587 |
| Heterotrimer | TYR 56.C - VAL 71.C | TYR 1274.C - VAL 1289.C | Intra-chain | 2 |
| Heterotrimer | TYR 56.A - VAL 71.A | TYR 1274.A - VAL 1289.A | Intra-chain | 2 |
| Heterotrimer | ILE 58.A - ILE 69.A | ILE 1276.A - ILE 1287.A | Intra-chain | 2 |
| Heterotrimer | TYR 59.B - VAL 74.B | TYR 1178.B - VAL 1193.B | Intra-chain | 1.99206 |
| Heterotrimer | ILE 61.B - ILE 72.B | ILE 1180.B - ILE 1191.B | Intra-chain | 1.99206 |
| Heterotrimer | ILE 58.C - ILE 69.C | ILE 1276.C - ILE 1287.C | Intra-chain | 1.99206 |
| Heterotrimer | ASP 46.B - THR 43.B | ASP 1165.B - THR 1162.B | Intra-chain | 1.96032 |
| Heterotrimer | ARG 39.C - PRO 60.C | ARG 1257.C - PRO 1278.C | Intra-chain | 1.94444 |
| Heterotrimer | ARG 42.B - PRO 63.B | ARG 1161.B - PRO 1182.B | Intra-chain | 1.88889 |
| Heterotrimer | CYS 41.C - THR 80.C | CYS 1259.C - THR 1298.C | Intra-chain | 1.69841 |
| Heterotrimer | ASP 59.C - GLN 62.C | ASP 1277.C - GLN 1280.C | Intra-chain | 1.57143 |
| Heterotrimer | CYS 41.A - THR 80.A | CYS 1259.A - THR 1298.A | Intra-chain | 1.53968 |
| Heterotrimer | ASP 43.A - CYS 47.A | ASP 1261.A - CYS 1265.A | Intra-chain | 1.5 |
| Heterotrimer | ASP 59.A - GLN 62.A | ASP 1277.A - GLN 1280.A | Intra-chain | 1.47619 |
| Heterotrimer | CYS 44.B - THR 83.B | CYS 1163.B - THR 1202.B | Intra-chain | 1.46032 |
| Heterotrimer | ARG 45.B - PHE 246.B | ARG 1164.B - PHE 1365.B | Intra-chain | 1.36508 |
| Heterotrimer | ARG 39.A - ASP 43.A | ARG 1257.A - ASP 1261.A | Intra-chain | 1.33333 |
| Heterotrimer | ASP 43.C - CYS 47.C | ASP 1261.C - CYS 1265.C | Intra-chain | 1.30159 |
| Heterotrimer | ASP 43.C - THR 40.C | ASP 1261.C - THR 1258.C | Intra-chain | 1.27778 |
| Heterotrimer | ASN 61.A - GLN 133.A | ASN 1279.A - GLN 1351.A | Intra-chain | 1.24603 |
| Heterotrimer | ASN 61.C - GLN 133.C | ASN 1279.C - GLN 1351.C | Intra-chain | 1.03968 |
| Heterotrimer | ASN 64.B - GLN 134.B | ASN 1183.B - GLN 1253.B | Intra-chain | 1.03175 |
| Heterotrimer | ASP 70.B - GLN 134.B | ASP 1189.B - GLN 1253.B | Intra-chain | 1 |
| Heterotrimer | ASP 67.A - GLN 133.A | ASP 1285.A - GLN 1351.A | Intra-chain | 0.992063 |
| Heterotrimer | ARG 42.C - LEU 246.C | ARG 1260.C - LEU 1464.C | Intra-chain | 0.984127 |
| Heterotrimer | ASN 61.A - ASP 67.A | ASN 1279.A - ASP 1285.A | Intra-chain | 0.960317 |
| Heterotrimer | ASN 61.C - ILE 132.C | ASN 1279.C - ILE 1350.C | Intra-chain | 0.928571 |
| Heterotrimer | ARG 45.B - LEU 49.B | ARG 1164.B - LEU 1168.B | Intra-chain | 0.920635 |
| Heterotrimer | ASN 64.B - ASP 70.B | ASN 1183.B - ASP 1189.B | Intra-chain | 0.84127 |
| Heterotrimer | ARG 39.C - ASP 43.C | ARG 1257.C - ASP 1261.C | Intra-chain | 0.801587 |
| Heterotrimer | ASN 64.B - THR 133.B | ASN 1183.B - THR 1252.B | Intra-chain | 0.690476 |
| Heterotrimer | ARG 42.C - MET 46.C | ARG 1260.C - MET 1264.C | Intra-chain | 0.674603 |
| Heterotrimer | ARG 42.A - MET 46.A | ARG 1260.A - MET 1264.A | Intra-chain | 0.65873 |
| Heterotrimer | ASN 61.A - ILE 132.A | ASN 1279.A - ILE 1350.A | Intra-chain | 0.515873 |
| Heterotrimer | ARG 42.B - ASP 46.B | ARG 1161.B - ASP 1165.B | Intra-chain | 0.253968 |
| Heterotrimer | ASP 67.C - GLN 133.C | ASP 1285.C - GLN 1351.C | Intra-chain | 0.222222 |
| Heterotrimer | ASN 61.C - ASP 67.C | ASN 1279.C - ASP 1285.C | Intra-chain | 0.142857 |

## Supplementary Figures

**Supplementary Figure S1.**
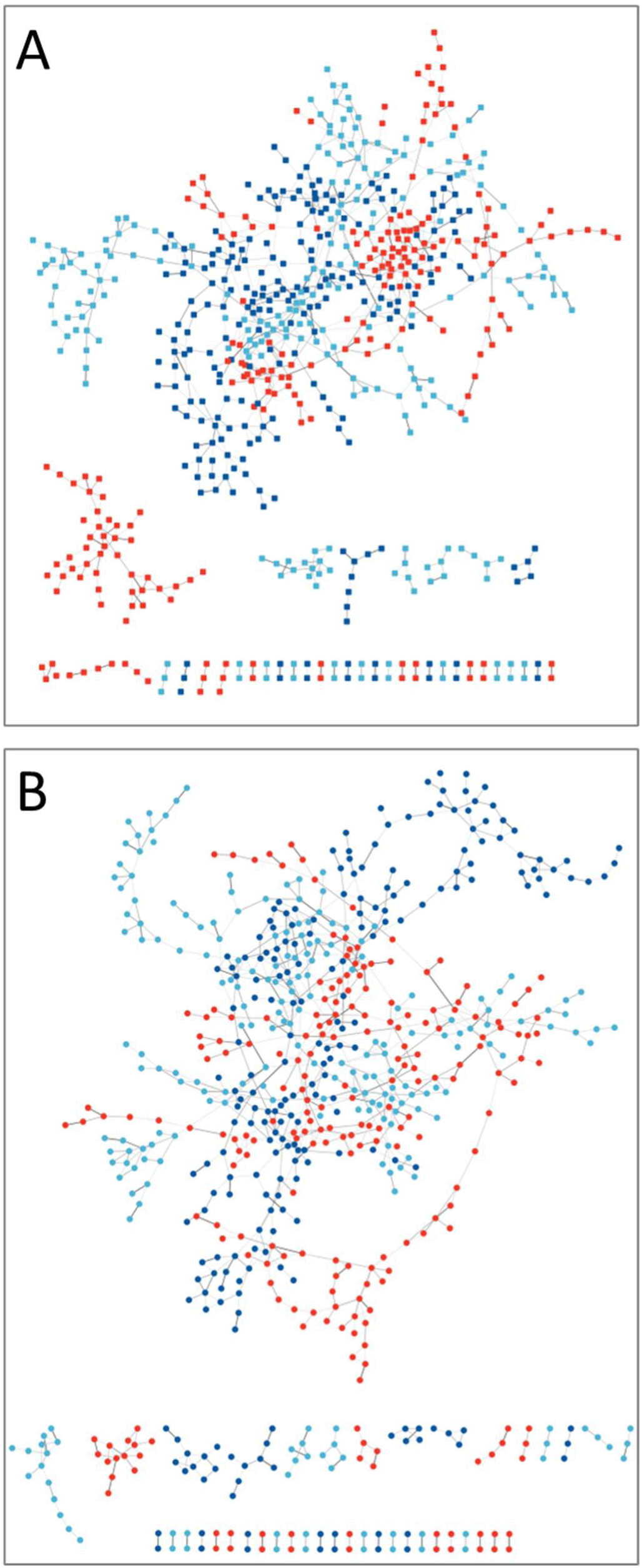
Cytoscape chimera full hydrogen bonding networks. A: Heterotrimer. B: Homotrimer. Red denotes the alpha-2 chain for the heterotrimer (A) and the alpha-1 (2) chain for the homotrimer (B). Blues represent the other alpha-1 chains in each trimer.

**Supplementary Figure S2.**
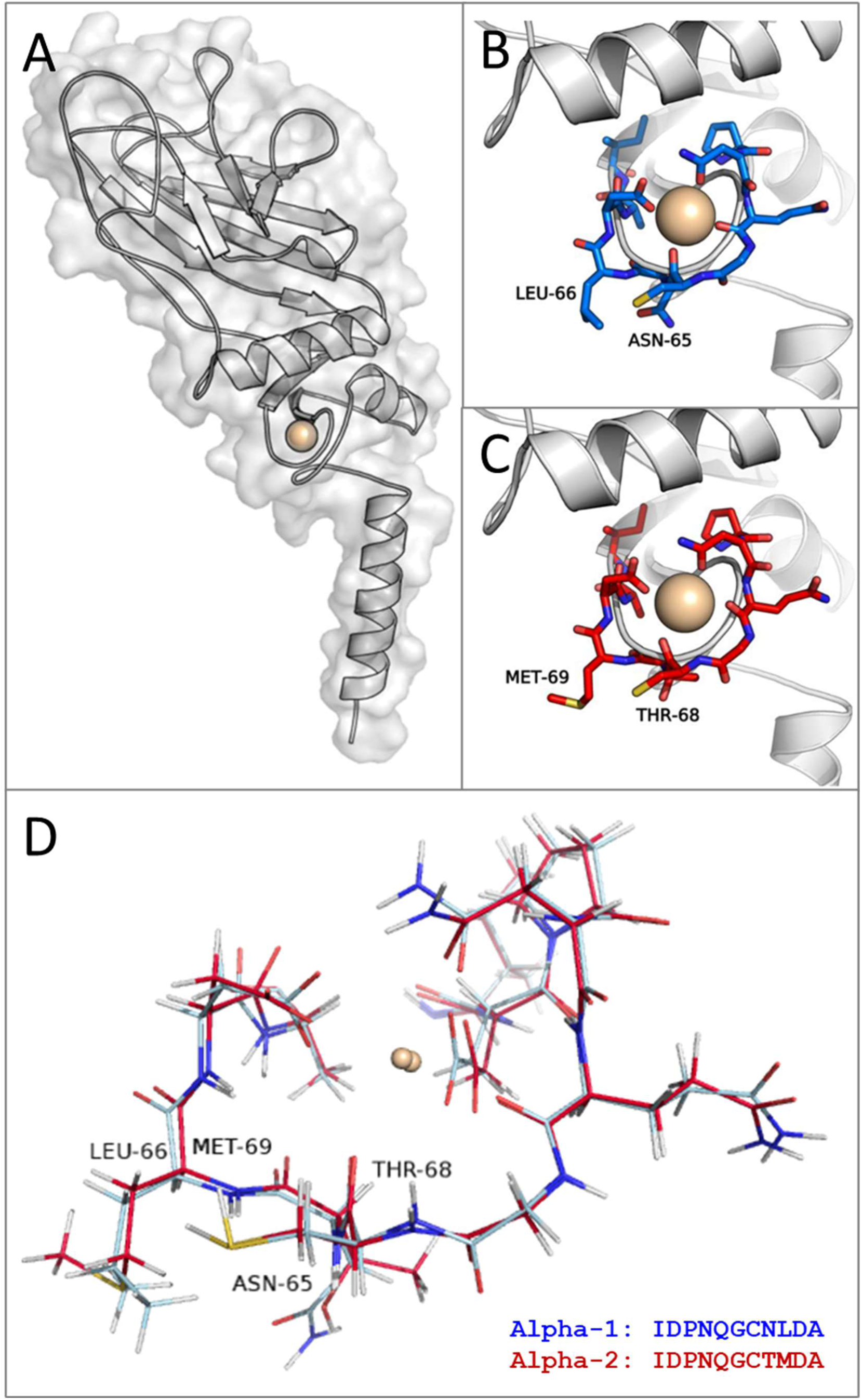
Calcium coordination loops in the alpha (I) chain monomers. A: Surface representation of an α1(I) monomer, demonstrating that calcium is exposed to solvent. B, C: Differences between the α1(I) (B), and α2(I) (C) calcium binding loops. D: Structure and sequence of the α1(I) (blue) and α2(I) (red) calcium-binding loops. Calcium ions are shown as spheres.

**Supplementary Figure S3.**
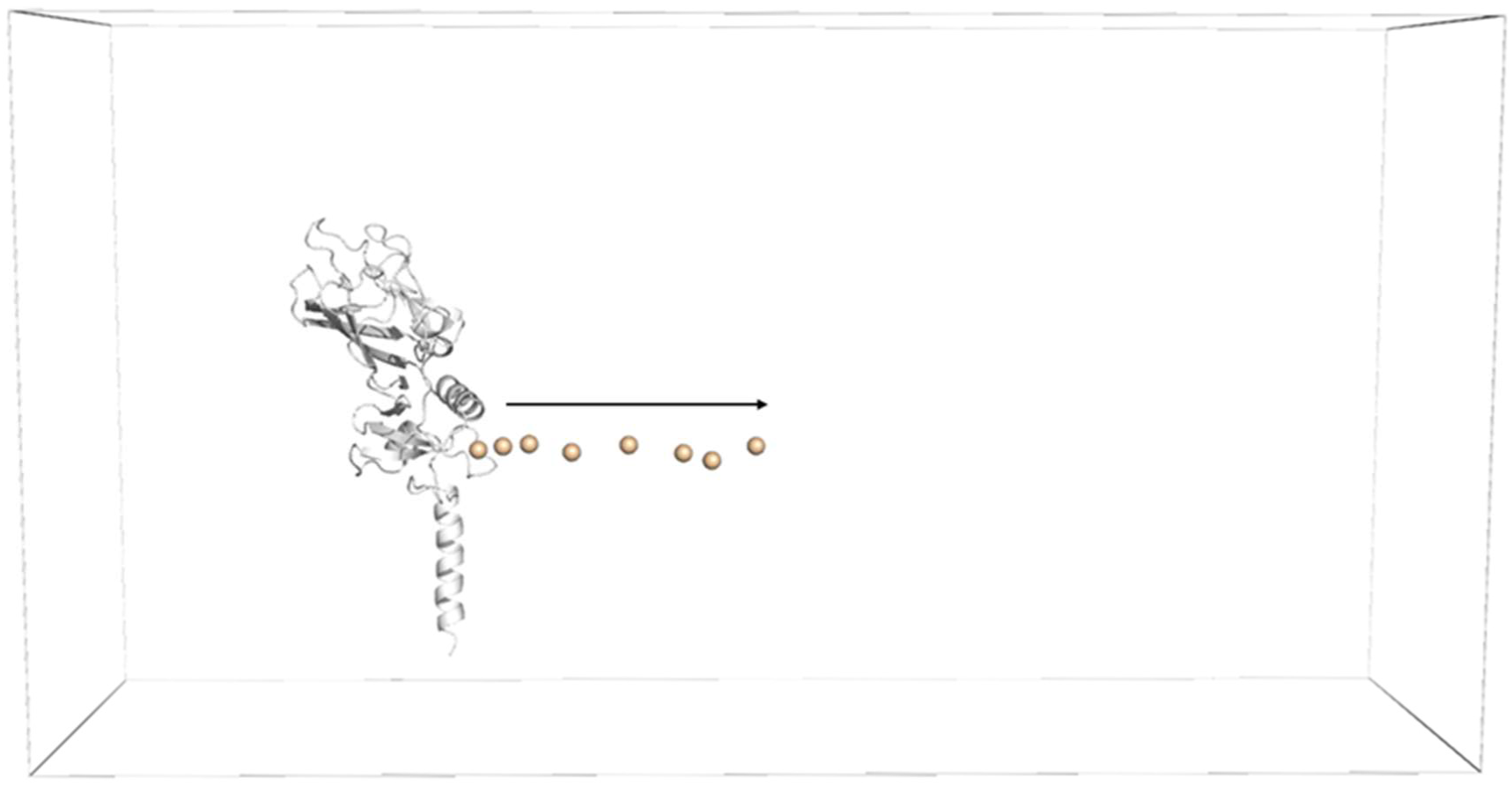
Centre of mass pulling simulation for a calcium ion extracted from its binding site in a monomer by steered molecular dynamics. An alpha-1 monomer is shown in white and the calcium ion as a wheat-coloured sphere. The box defines the periodic boundary conditions. Multiple frames of the calcium moving away are shown along the arrow, which was the direction of the pull force.

**Supplementary Figure S4.**
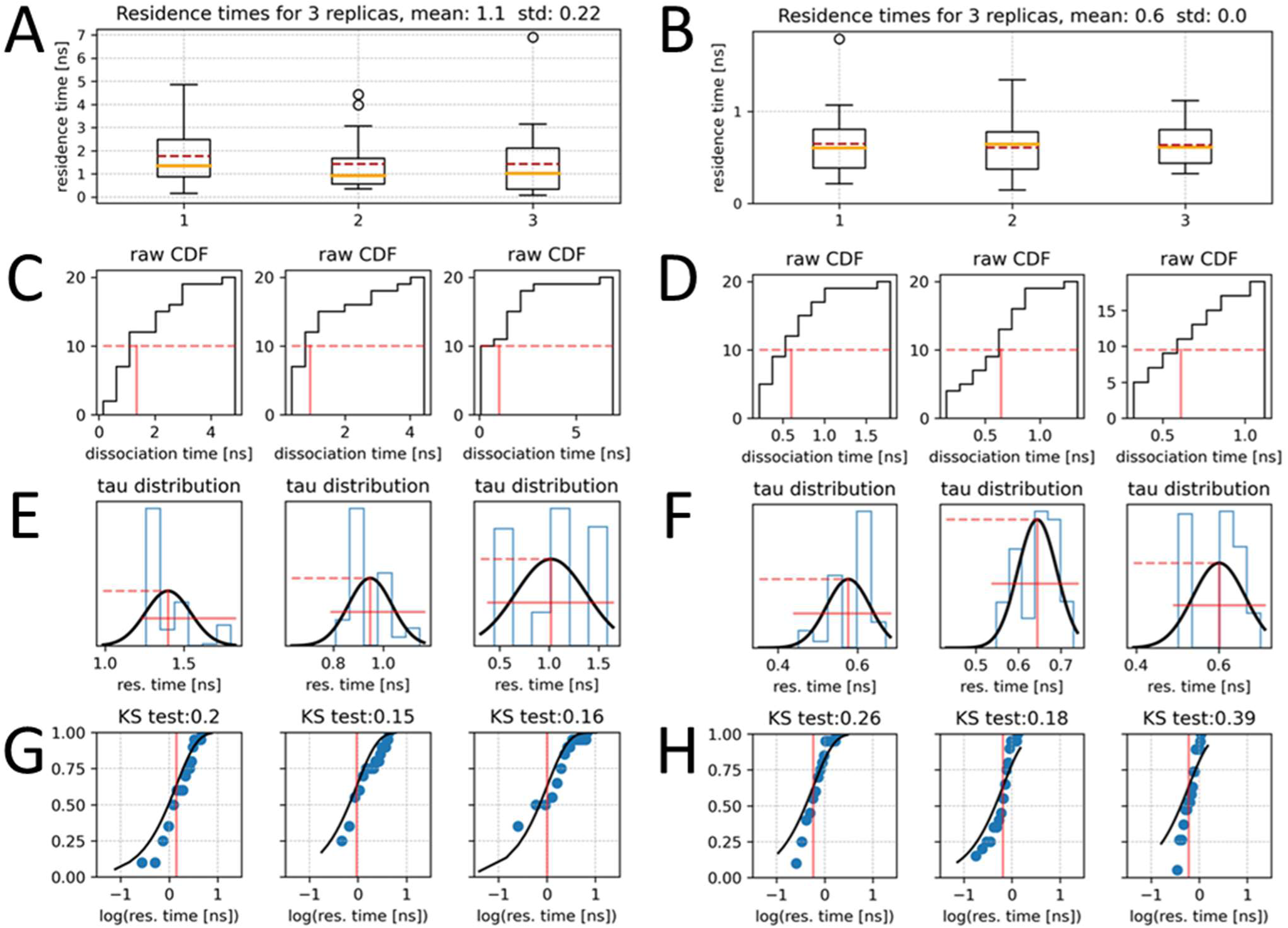
τRAMD residence times and statistical analysis for the α1(I) and α2(I) chains. A, B: The residence times for 3 replicas for the alpha-1(I) (A) and alpha-2(I) chain (B). Box plots represent 25 τRAMD results. The median value is shown as an orange line and the mean value as a red dashed line, C, D: Analysis of the time at which 50% of the trajectories had dissociated for the alpha-1(I) (C) and alpha-2(I) chain (D). E, F: Fit of a normal distribution to the data for the alpha-1(I) (E) and alpha-2(I) chain (F). G, H: Kolmogorov–Smirnov test results for the for the alpha-1(I) (G) and alpha-2(I) chain (H). The line is the Poisson cumulative distribution function and the blue dots the cumulative density function (CDF).

**Supplementary Figure S5.**
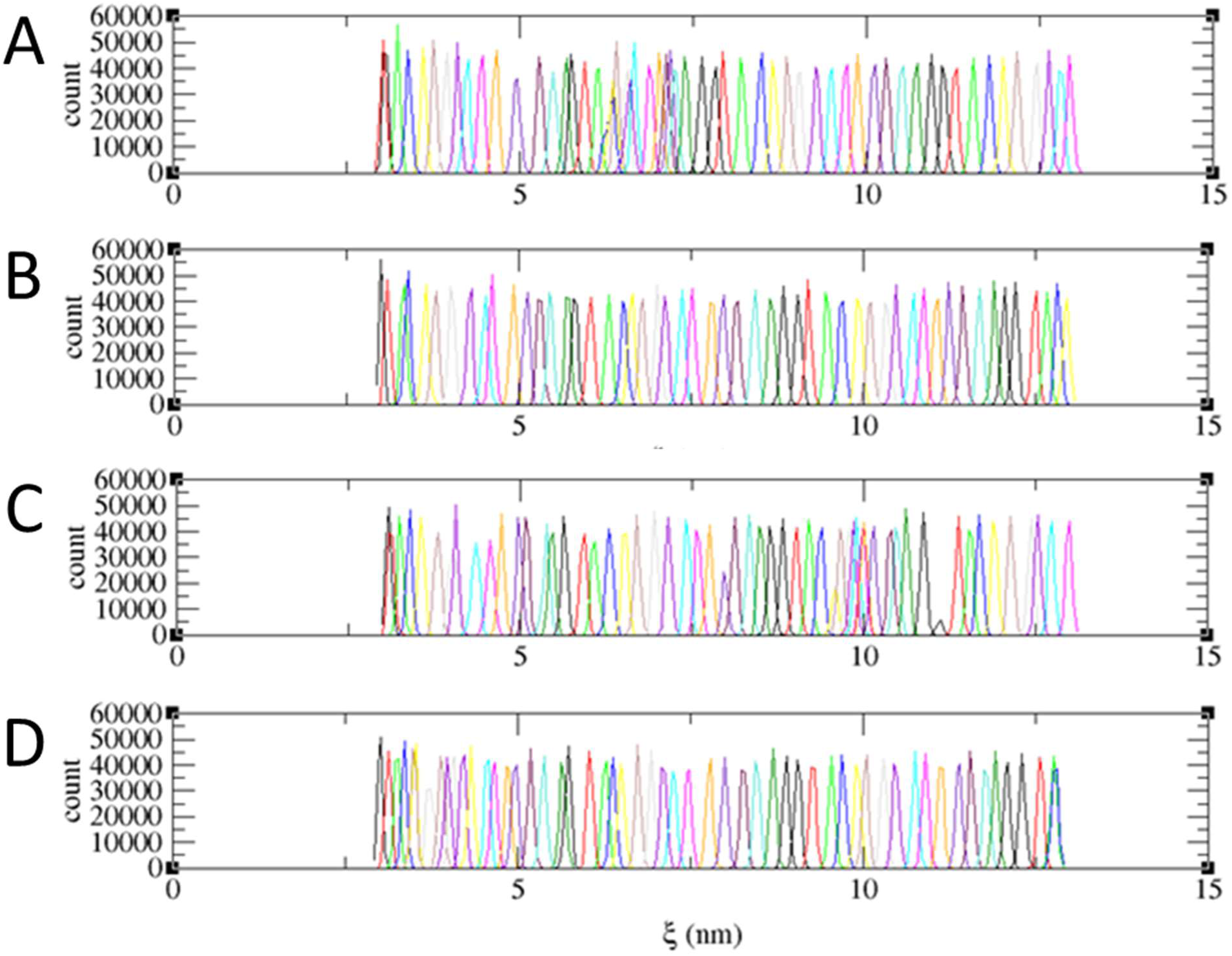
Histograms of the sampling windows for trimer pulling simulations. Each bin represents a sampling window. The overlap between the bins demonstrates good sampling of the system. A: Heterotrimer histogram, B: Apo-heterotrimer histogram, C: Homotrimer histogram, D: Apo-homotrimer histogram.

**Supplementary Figure S6.**
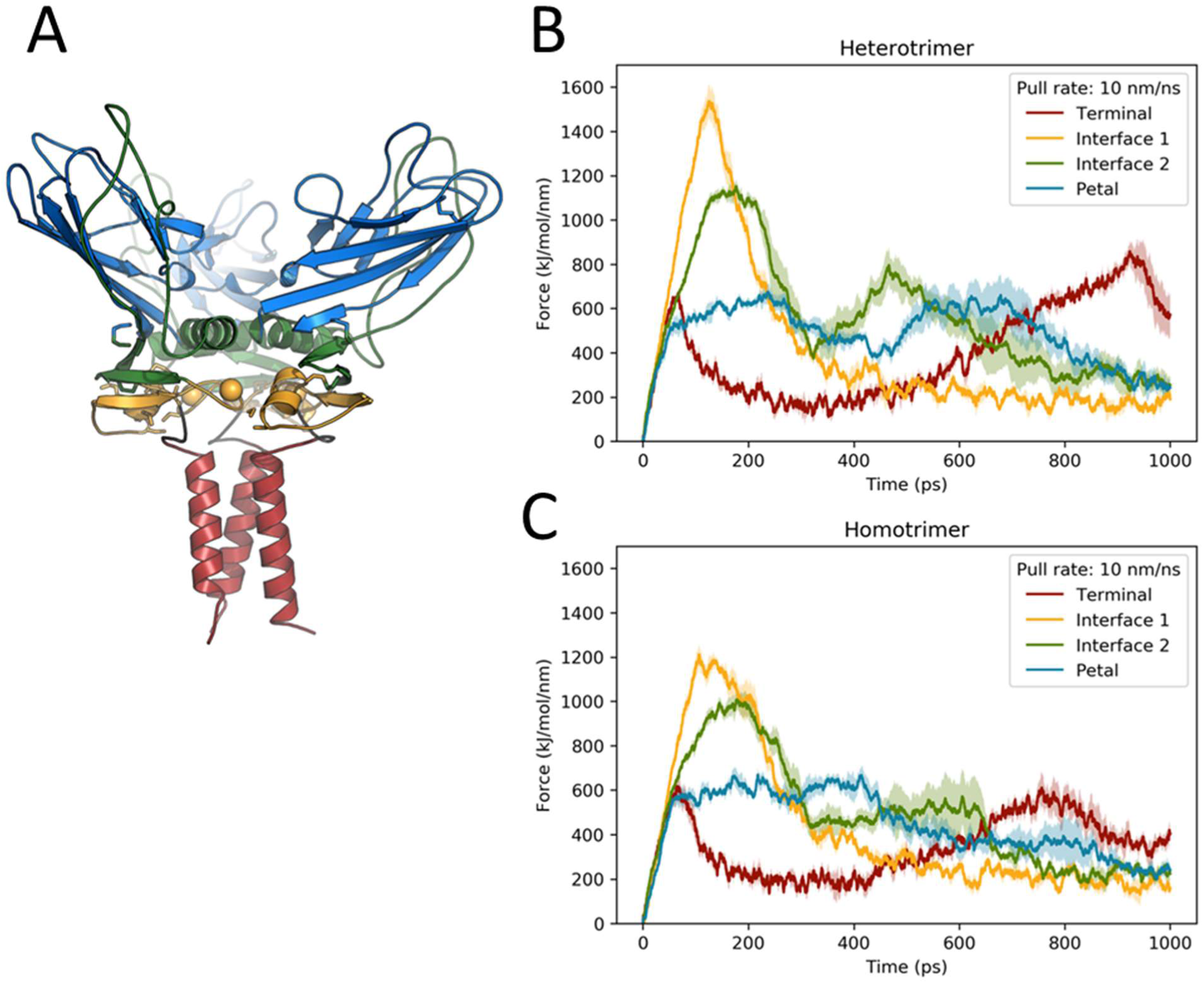
Pull group optimisation for trimer pulling simulations. The trimer was divided into different regions from which the pulling then took place. Red: terminal region. Yellow: calcium-binding region (interface 1). Green: central alpha-helices, including a loop that extends into the petal (interface 2). Blue: petal region. A: Trimer with terminal (red), interface regions 1 (yellow) and 2 (green) and petal (blue) regions distinguished by colour. B, C: Force-time curves for each region of the heterotrimer (B) and homotrimer (C).

**Supplementary Figure S7.**
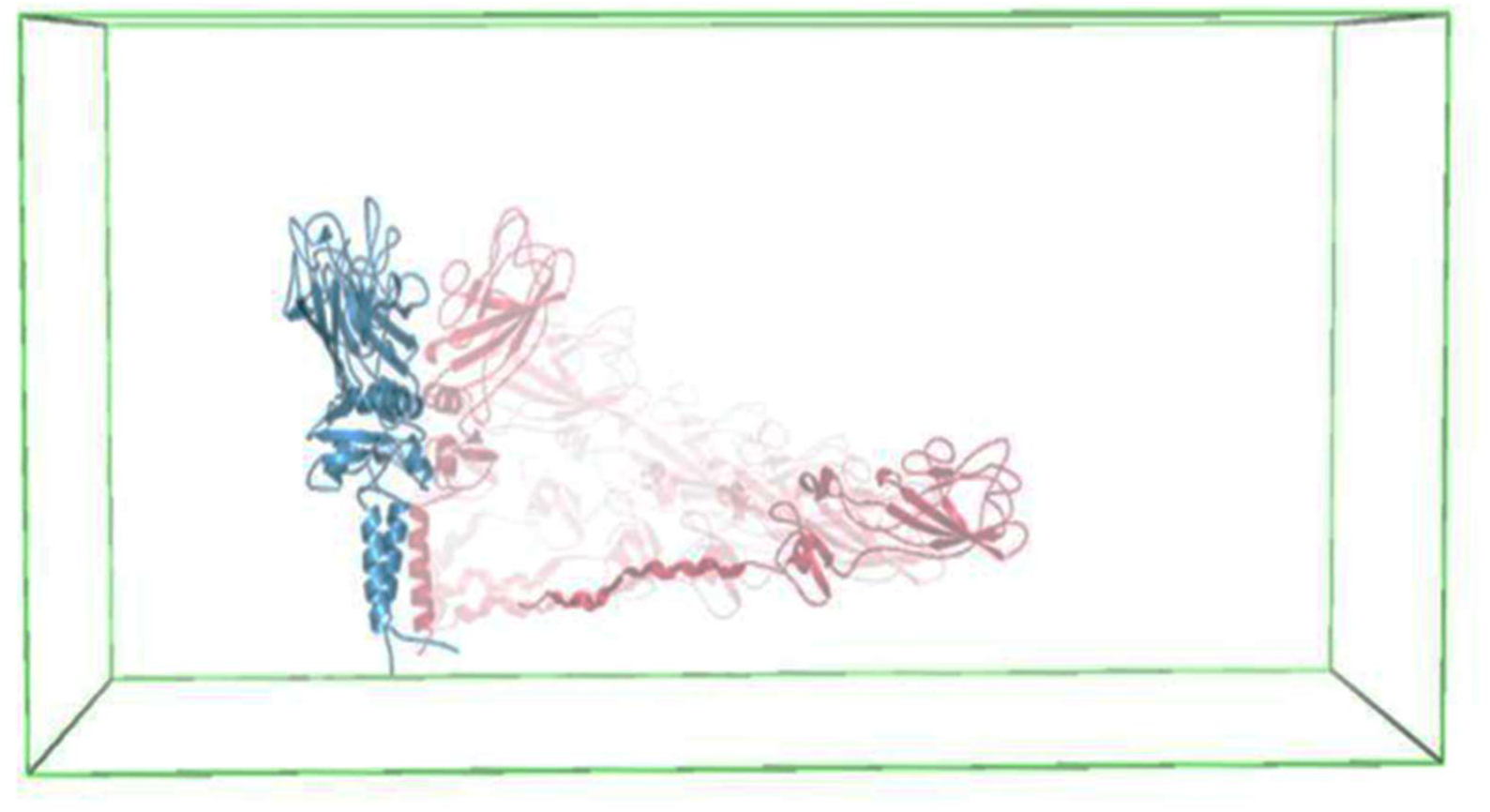
Sample unbinding pathway for the heterotrimer. An α2(I) chain is dissociating from two α1(I) chains along the collective variable (z-axis). The end configuration is shown as the darkest shade of red; the earliest configuration is shown as the next darkest shade. The α1(I) chains are shown in blue.

